# Determinants of spring migration departure dates in a New World sparrow: weather variables reign supreme

**DOI:** 10.1101/2023.11.17.567563

**Authors:** Allison J. Byrd, Katherine M. Talbott, Tara M. Smiley, Taylor B. Verrett, Michael S. Gross, Michelle L. Hladik, Ellen D. Ketterson, Daniel J. Becker

## Abstract

Numerous factors influence the timing of spring migration in birds, yet the relative importance of intrinsic and extrinsic variables on migration initiation remains unclear. To test for interactions among weather, migration distance, parasitism, and physiology in determining spring departure date, we used Dark-eyed Juncos (*Junco hyemalis hyemalis*) as a model migratory species known to harbor diverse and common haemosporidian parasites. Prior to spring migration departure from their wintering grounds in Indiana, USA, we quantified the intrinsic variables of fat, body condition (i.e., mass∼tarsus residuals), physiological stress (i.e., ratio of heterophils to lymphocytes), cellular immunity (i.e., leukocyte composition and total count), migration distance (i.e., distance to the breeding grounds) using stable isotopes of hydrogen from feathers, and haemosporidian parasite intensity. We then attached nanotags to determine the timing of spring migration departure date using the Motus Wildlife Tracking System. We used additive Cox proportional hazard mixed models to test how risk of spring migratory departure was predicted by the combined intrinsic measures, along with meteorological predictors on the evening of departure (i.e., average wind speed and direction, relative humidity, and temperature). Model comparisons found that the best predictor of spring departure date was average nightly wind direction and a principal component combining relative humidity and temperature. Juncos were more likely to depart for spring migration on nights with largely southwestern winds and on warmer and drier evenings (relative to cooler and more humid evenings). Our results indicate that weather conditions at take-off are more critical to departure decisions than the measured physiological and parasitism variables.

## Introduction

Over-wintering duration and timing of departure for avian spring migration is a process influenced by a host of extrinsic and intrinsic factors (Covino et al., 2015; Hurlbert and Liang, 2012; Satterfield et al., 2018). The timing of spring migration must balance the advantages of early arrival on the breeding grounds, including higher quality mates and territories (Gunnarsson et al., 2006; Møller, 1994; Newton, 2010; Rotics et al., 2018; Smith and Moore, 2005), with the risk of arriving before adequate food resources are available (Gunnarsson et al., 2006; Møller, 1994; Newton, 2010; Rotics et al., 2018; Smith and Moore, 2005). Stressors experienced on the wintering grounds can also affect the timing and duration of spring migration. Extrinsic factors such as changing weather conditions (Marra et al., 2005) and intrinsic factors such as high parasite intensity (Dietsch, 2005; Reed et al., 2003) can alter stopover duration as birds recover from or accommodate these additional physiological burdens (Schmaljohann et al. 2022; Skrip et al. 2015; Linscott and Senner 2021). The relative contribution of extrinsic and intrinsic factors experienced at the wintering grounds in shaping spring departure remains less well understood, and studies addressing these interactions are especially needed as anthropogenic pressures continue to influence the climate and natural habitat (Kubelka et al., 2022; Visser et al., 2009; Wilcove and Wikelski, 2008).

For decades, ornithologists have known that weather variables influence avian migration timing in both spring and fall (see:(Richardson, 1978). Factors that influence the probability of migratory departure include wind speed (Chapman et al., 2016; Drake et al., 2014; Kemp et al., 2010; Liechti, 2006; Nussbaumer et al., 2022), wind direction (Covino et al., 2015; Hebrard, 1971; Horton et al., 2016; Kemp et al., 2010; Lack, 1963; Lack and Eastwood, 1962; Sinelschikova et al., 2007), temperature (Hüppop and Winkel, 2006; Marra et al., 2005; Saino et al., 2007; Tøttrup et al., 2010; Usui et al., 2017), and relative humidity (Klaassen et al., 2012; Liechti, 2006; Schmaljohann et al., 2009; Serra-Cobo et al., 1998; Zhang and Wu, 2018). The high correlation among these weather variables adds to the difficulty in disentangling the influence of these factors from potential additional stressors. For example, warm air can hold more moisture than cold air, thus at the same absolute humidity, cooler air (perhaps counterintuitively) has higher relative humidity than warmer air (Lawrence, 2005).

The relationship between wind and bird migration has been studies observed for decades, and prevailing winds were a likely a selection factor in that shaping the evolution of migratory patterns (Alerstam 1979; Alerstam 1979; Evans 2009; Able 1972; Richardson 1978). Birds that are highly selective of favorable winds can maximize flight speed and minimize energy expenditure across both long and short-distance migratory flights (Alerstam 1979). Migrants have faster ground and air-speeds in spring migration than fall (Horton et al. 2016), and spring migration is typically completed over a shorter duration than fall (Newton 2010; La Sorte et al. 2013; La Sorte et al. 2016; Nilsson et al. 2013). As such, models that forecast bird migration rely heavily on wind data (Erni et al., 2002; Van Doren and Horton, 2018).

Intrinsic factors have also been recognized to affect readiness for migratory departure. Given the energetic costs of migration (Wikelski et al., 2003), adequate fat stores are integral in providing fuel for long-distance flight and larger stores can be a reliable predictor of migratory readiness(Price, 2010; Ramenofsky, 1990; Weber et al., 1994; Witter and Cuthill, 1993); correspondingly, measures of body condition (i.e., mass∼tarsus residuals) allow for standardized body size comparisons among individuals (Labocha and Hayes, 2012). The degree of physiological stress and immune state of individuals at the wintering grounds can also affect spring migratory timing. Preparation for long-distance migration increases plasma concentrations of corticosterone (the primary avian glucocorticoid responsible for maintaining homeostasis) in many songbirds (Holberton, 1999; Landys et al., 2004), which can facilitate earlier departure from the wintering grounds or spring stopover (Eikenaar et al., 2013, 2017). Elevated corticosterone can shift the composition of leukocytes in blood, subsequently elevating the ratio of heterophils to lymphocytes (HL ratios; Davis et al., 2008). While plasma corticosterone levels are highly sensitive to the acute stress of capture, stress-induced changes in leukocyte profiles occur more slowly, such that HL ratios can serve as a more tractable approximation of energetic costs prior to migration (Davis and Maney, 2018). In a similar fashion, the energetic cost of migratory preparation can induce trade-offs with the immune system (Lochmiller and Deerenberg, 2000), such that migrants may downregulate immune activity or rely primarily on less costly immune defenses. For example, several thrush species show lower total leukocyte counts upon arrival at stopover sites in spring, reflecting such trade-offs (Owen and Moore, 2008). Variation in immune investment may in turn affect departure decisions; recent work demonstrated that songbirds with higher titers of natural antibodies and immunoglobulin Y have longer spring stopovers (Brust et al., 2022).

Related to seasonal variation in physiological stress and immunity, songbirds may also experience additional overwintering stressors that affect spring migration. For example, exposure to toxins such as neonicotinoids in wintering food sources (Eng et al., 2017; Goulson, 2014) can delay migration departure or increase stopover duration as birds recover from or accommodate these burdens. An additional such stressor could be through parasite infection, such as that from dipteran-borne haemosporidian blood parasites (i.e., the genera *Plasmodium*, *Haemoproteus*, and *Leucocytozoon*). Experimental infections of captive birds have shown migration-relevant physiological costs of infection, such as reduced general activity (Mukhin et al., 2016; Yorinks and Atkinson, 2000) and body condition (Atkinson et al., 1995) as well as shifts in the development of migratory restlessness (Kelly et al., 2016, 2020). Observational studies have also identified impacts of haemosporidian infection on body condition in migrating birds (Garvin et al., 2006; Merrill et al., 2018), on the timing of arrival to the breeding grounds (Asghar et al., 2011; Santiago-Alarcon et al., 2013), and on autumn departure timing (Ágh et al., 2019). However, it remains unclear how haemosporidian infections interact with extrinsic and other intrinsic factors to affect migration timing.

In this study, we used overwintering migratory Dark-eyed Juncos (*Junco hyemalis hyemalis;* hereafter “junco”) to test the hypothesis that physiological state and haemosporidian infection interact with weather conditions and migration distance to shape spring migratory departure timing. Haemosporidia have been well characterized in Dark-eyed juncos, with chronic (i.e., long-term) infections detected in wintering and breeding populations (Becker et al. 2020; Slowinski et al. 2018; Talbott et al. 2022; Martínez-Renau et al. 2022; Becker et al. 2019; Deviche, Greiner, and Manteca 2001; Ferrer 2022). We used stable isotopes of hydrogen from feathers to estimate the distance to breeding grounds for each individual (Bowen et al., 2014; Hobson, 1999; Hobson et al., 2012; Rubenstein and Hobson, 2004; Wunder, 2010) and the Motus Wildlife Tracking System to determine departure date (Taylor et al., 2017). We predicted that birds with higher fat reserves, low intensity or no haemosporidian parasitism, lower HL ratios, and lower total leukocyte counts would depart earlier in the year and during evenings of following winds (i.e., tailwind). We also predicted that wind direction (following winds) would be more important than wind speed, and that headwinds would be the strongest deterrent to departure, even more so than intrinsic predictors.

## Methods

### Junco sampling and captive housing

We caught wild juncos from 31 January to 26 February 2020 at four locations within 15 km of Bloomington, Indiana. None of the birds were captured at the release site, and we systematically rotated capture efforts through all four sites to avoid confounding results by capture date or location. We captured juncos using baited mist nets and walk-in traps, took standard morphometric measurements including mass and body fat (Fudickar et al., 2016; Jawor et al., 2006; Singh et al., 2019), and fitted each bird with an aluminum US Fish & Wildlife Service leg band (federal permit # 20261, Indiana state permit 20-528).

While capture efforts were ongoing, juncos were held at an indoor aviary at Indiana University to prevent additional exposure to arthropod vectors of haemosporidia. Juncos were provided *ad libitum* water and food (mealworms, organic millet, organic sunflower seeds, and a blended mixture of organic millet, organic carrots, and organic blueberries) and could fly freely in 6.4 x 3.2 m rooms. Light cycles for rooms reflected the natural photoperiod of Bloomington, Indiana at the time of the study.

As a potential additional extrinsic factor, juncos in this study were randomly dosed with either a neonicotinoid (imidacloprid suspended in sunflower seed oil) or control (sunflower seed oil) to experimentally test impacts on migration timing. However, imidacloprid metabolites of 5-OH-imidacloprid and imidacloprid-olefin were detected in only one bird’s post-dose plasma at 46.67 ng/mL and 72.23 ng/mL. This individual was removed from all subsequent analyses. Imidacloprid or metabolites were not detected in pre-dose or post-dose (6 hours after) plasma of any other individuals (*n* = 37; Supp. Appx A, Table S1). Therefore, we did not include imidacloprid exposure as a possible predictor of migratory timing. Possible explanations for the absence of imidacloprid detection in plasma samples are that the dose was regurgitated or rapidly metabolized by all birds.

On the days of release (3 and 4 March 2020), we attached a nanotag to each bird (Lotek model NTQB2-2, 11 x 5 x 4 mm, 0.32g, ≦ 0.5 g total mass including leg harness). Birds were released at Kent Farm Research Station (located nine miles east of Bloomington, IN), an area frequented by overwintering juncos; we provided organic seed in feeders and on the ground in an attempt to minimize departure from the area due to resource limitation (Bridge et al., 2010) .

### Hematological analyses

At capture, we collected ≤ 150 μL blood from each bird by pricking the brachial vein with a sterile needle, followed by collection with heparinized capillary tubes. We also collected the first secondary feather from each bird for stable isotope analysis (Fudickar et al., 2016). We separated plasma and red blood cells (centrifuged for 10 min at 10,000 rpm) and stored samples at -20℃ until DNA extraction using a Maxwell RSC Whole Blood DNA Kit (Promega). Any birds unable to be sexed by wing length and plumage were verified using sexing PCR of extracted DNA (Griffiths et al., 1998). Only males (*n*=37) were included to control for sex differences in migration timing, as female juncos migrate earlier than males (Nolan and Ketterson, 1990).

Because birds were held in captivity for variable lengths between capture and release with nanotags (x̄ =25, range of 13 to 33 days), we collected additional blood samples 24 hours prior to release, using the blood collection protocol described above. We prepared thin blood smears on glass slides stained with Wright–Giemsa (Quick III, Astral Diagnostics). We then evaluated leucocyte profiles and haemosporidian intensity using light microscopy (AmScope, B120C-E1). A single observer (TBV) recorded the number of leukocytes under 400X magnification across 10 random fields to estimate inflammatory state and investment in cellular immunity. A differential count was then performed by recording the identity of the first 100 leukocytes (heterophils, lymphocytes, monocytes, eosinophils, and basophils) at 1000X magnification (oil immersion;(Campbell, 1995). We then screened 100 fields of view at 1000X magnification for *Plasmodium*, *Haemoproteus*, and *Leucocytozoon (Becker et al., 2019, 2020; Cosgrove, 2005; Valkiunas et al., 2008)*. We derived the mean total leukocyte count, HL ratios, and the total number of haemosporidian-infected erythrocytes (parasite intensity) as three predictor variables. Usable blood smears were available for 34 of our 37 tagged individuals.

### Hydrogen isotope analysis

We used stable isotopic analysis of hydrogen (δ^2^H) in feathers to infer probable breeding latitude and median distance from capture point(Bowen et al., 2014; Hobson, 1999; Hobson et al., 2012; Rubenstein and Hobson, 2004; Wunder, 2010). Feathers were cleaned using a 2:1 chloroform:methanol solution to remove external oils and contaminants. We used a forced isotopic equilibration procedure to ensure exchangeable hydrogen with a water vapor of known isotopic composition in a flow-through chamber system at 115°C (Sauer et al., 2009; Schimmelmann, 1991). Samples were analyzed using a thermal conversion element analyzer coupled with a ThermoFinnigan Delta Plus XP isotope ratio mass spectrometer at the Indiana University Stable Isotope Research Facility. Isotopic data are reported in standard per mil notation (‰) relative to VSMOW (Vienna Standard Mean Oceanic Water) using two reference materials: USGS77 (polyethylene powder) and hexatriacontane 2 (C_36_ n-alkane 2). Analytical precision was ± 1.0‰ for δ^2^H values. We calculated the isotopic composition of the non-exchangeable hydrogen per sample, assuming a 17% exchangeability rate for feathers (Schimmelmann, 1991; Schimmelmann et al., 1999).

We performed Bayesian geographic assignments for the breeding location of individual birds based on feather δ^2^H values using the *assignR* package (Ma et al., 2020). To calibrate our precipitation-feather isoscape, we used growing season precipitation isoscape rasters from waterisotopes.org (Bowen et al., 2005; Bowen and Revenaugh, 2003) and the isotopic composition of non-migratory Dark-eyed juncos from previous studies(Becker et al., 2019; Hobson et al., 2012). Median latitudinal and distance estimates were calculated from geographic assignment probability maps by extracting the geographic cell coordinates with the highest posterior probability (top 10%) using the *raster* package (Hijmans, 2019). Migration distance was defined as the distance from the release site to the centroid of the estimated breeding polygon (Becker et al., 2022; Wanamaker et al., 2020).

### Motus data processing

Motus data (project 240; available at https://motus.org/data) were downloaded on 25 July 2020, four months after the known period of spring migration in Indiana of wintering juncos (Ketterson and Nolan, 1982). Data were filtered and cleaned following a standard protocol using the *motus* package in R(Taylor et al., 2017). Specifically, we removed runs with a low probability of being a true detection (i.e., motusFilter=0) and adopted a moderately strict filter to be more conservative about detection inclusions. We then derived the departure date as the last day (ordinal date) a bird was detected at the Kent Farm Research Station Motus station. We additionally report cases in which birds were detected at other Motus stations between banding and our data download date.

### Weather data

We obtained relative humidity data from ncei.noaa.gov (Diamond et al., 2013) and all remaining weather variables (humidity, wind speed, wind direction [converted to wind rose] and weather type) from timeanddate.com (Thorsen, 1995-2023). We averaged hourly data from 1800-2400 on departure date evenings (thus, variables are labeled average humidity and so on).

Because average relative humidity and average temperature are dependent (Lawrence, 2005), we conducted a principal components analysis (PCA) of these two variables, with variables centered and scaled to have unit variance. The first PC (hereafter weather PC1) explained 56% of the variation and was loaded negatively by average relative humidity (–0.71) and positively by average temperature (0.71); resulting values indicate increasingly warm and dry evening weather. This variable was used in downstream analyses alongside average wind speed and average wind direction.

### Statistical analyses

Prior to analyses of migration timing, we used a linear mixed effects model with a random effect of site to derive an index of body condition through the residuals of a regression of mass (measured prior to release) on tarsus length (Schulte-Hostedde et al., 2005; Wanamaker et al., 2020).

We modeled days until spring migratory departure under an event-time analysis framework, using additive Cox proportional hazard mixed models (CPHMMs, (Therneau and Grambsch, 2000) and Grambsch, 2000). These semi-parametric models allow determining how the risk of departure changes with covariates that can also vary with time, and such models have been applied to study migration timing in both avian and non-avian systems (Castro-Santos and Haro, 2003; Dossman et al., 2015). We used the *mgcv* package to fit CPHMMs with a random effect of site. To test the direct and interactive effects of extrinsic (i.e., weather PC1, wind direction, wind speed) and intrinsic predictors (i.e., fat, body condition, total leukocytes, HL ratios, haemosporidian intensity) on risk of spring departure (Wood, 2017).

Including all predictor variables and biologically relevant interactions in a single full model was not possible given our sample size (*n* = 34, excluding birds without hematology data). We instead built 10 candidate CPHMMs representing *a priori* hypotheses of additive and interactive effects while restricting models to at most three fixed effects to limit overfitting (Burnham and Anderson, 2002). Most predictors showed low collinearity (ρ ranged from –0.63 to 0.62, x̄ = 0.01), and moderately correlated predictors (i.e., weather PC1 and wind speed, ρ = –0.63; body fat and condition, ρ = 0.62) were excluded from the same models. All predictors were modeled using thin plate splines with smoothing penalty with the exception of wind direction, which used a cyclic cubic spline to account for circular data. Interaction terms were modeled as tensor products. Our models represented additive effects of intrinsic variables only (e.g., fat, haemosporidian intensity, and total leukocytes), additive effects of extrinsic variables only (e.g., wind direction and weather PC1), additive effects of both intrinsic and extrinsic variables (e.g., haemosporidian intensity, HL ratios, and weather PC1), interactive effects of intrinsic variables (e.g., effects of haemosporidian intensity dependent on HL ratios), interactive effects of extrinsic variables (e.g., effects of wind speed depend on wind direction), and interactive effects of intrinsic and extrinsic variables (e.g., effects of weather PC1 depend on haemosporidian intensity). We compared CPHMMs fit with maximum likelihood using Akaike information criterion adjusted for small sample size (AICc) with the *MuMIn* package(Bartoń, 2013). We also derived Akaike weights (*w_i_*) to facilitate comparison and considered models within two ΔAICc of the top model to be competitive. Competitive models were refitted to the full dataset, with CPHMM predictions visualized as relative hazards with 95% confidence intervals (Nakagawa and Schielzeth, 2013).

## Results

### Hydrogen isotopes and likely breeding origins

Hydrogen isotopic composition of junco feathers ranged from -158.9 to -95.8%, reflecting median breeding locations from 66 to 51°N, respectively (Fig. 1). Maximum and minimum latitudinal estimates spanned from 70 to 36 °N. Estimated distances to breeding grounds accordingly ranged from approximately 2086 to 3900 kilometers (x̄ =3297, SE=78.4). Incidentally, we detected six of our 37 tagged juncos at other Motus towers following their departure from Indiana (Fig. 1). Five birds were detected at the Lake Petite station in Wisconsin (42.5117°, -88.5488°) and one bird was detected at the Werden station (42.7551°, -80.2724°) in Ontario, Canada.

**Figure 1.**
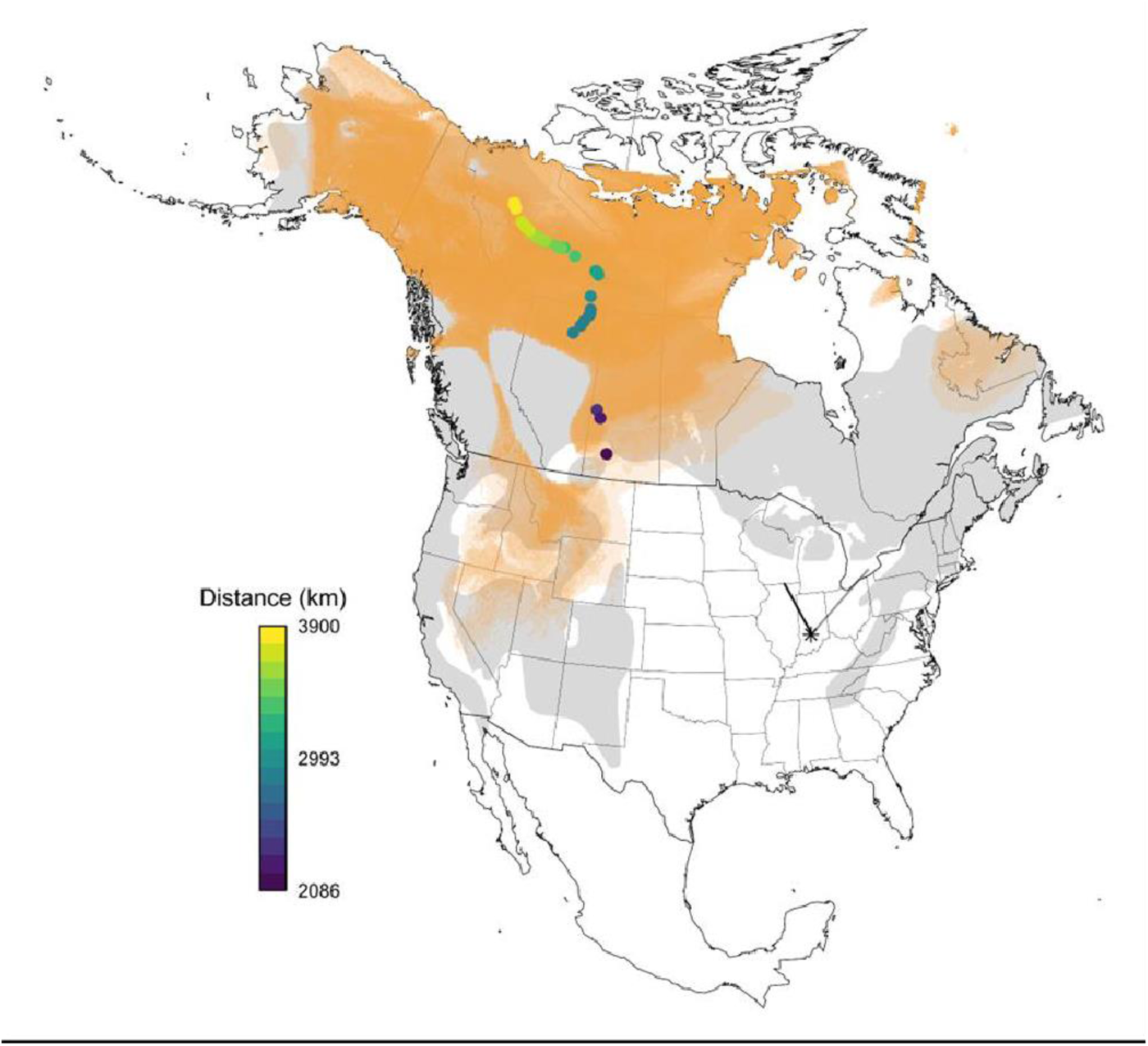
Estimated breeding locations and migratory distances based on junco feather δ^2^H. Orange shading represents overlapping geographic breeding assignments of individuals. Estimates reflect cells within the top 10% highest posterior probability (Bowen et al., 2014; Ma et al., 2020; Wunder, 2010). Points on the map represent the centroid of each estimated breeding polygon, colored by distance to location of capture (star; Bloomington, IN). Lines radiating from the star represent subsequent Motus detections after departure. The seasonal breeding range for *Junco hyemalis(Baillie et al., 2004)* is shown in light grey (the species’ full range map was used to construct geographic assignment maps).

### Haemosporidian infection

We identified haemosporidian infections in 11 of 34 juncos prior to release (32.4%, 95% CI: 19.1– 49.2%). Only two individuals had detectable *Haemoproteus* spp. infection (intensity=16–23 infected erythrocytes from 100 fields of view), and nine individuals had detectable *Leucocytozoon* spp. infection (intensity=1–7 infected erythrocytes from 100 fields of view); no individuals harbored co-infecting haemosporidian parasites. We also opportunistically detected *Trypanosoma* spp. in a single bird (1/34, two parasites were detected from 100 fields of view), and this individual was not infected by either *Haemoproteus* spp. nor *Leucocytozoon* spp. parasites.

### Extrinsic and intrinsic predictors of migration timing

All birds departed 8–31 days after release (x̄ =19.11 ± 0.81 SE), between March 11 and April 3 2020 (Fig. 2). Comparison among 10 CPHMMs predicting risk of departure date as a function of extrinsic and intrinsic factors identified only one competitive model, which included the nonlinear additive effects of wind speed and weather PC1 (*w_i_*=0.61; Table 1). Wind direction had the greatest relative importance among predictors (99%), followed by weather PC1 (62%); wind speed, the interaction between wind speed and weather PC1, and estimated migration distance all had lesser importance (38%, 22%, and 16%, respectively). All intrinsic predictors (i.e., fat, body condition, total leukocytes, HL ratios) and their interactions were unimportant (≤ 1%).

**Figure 2.**
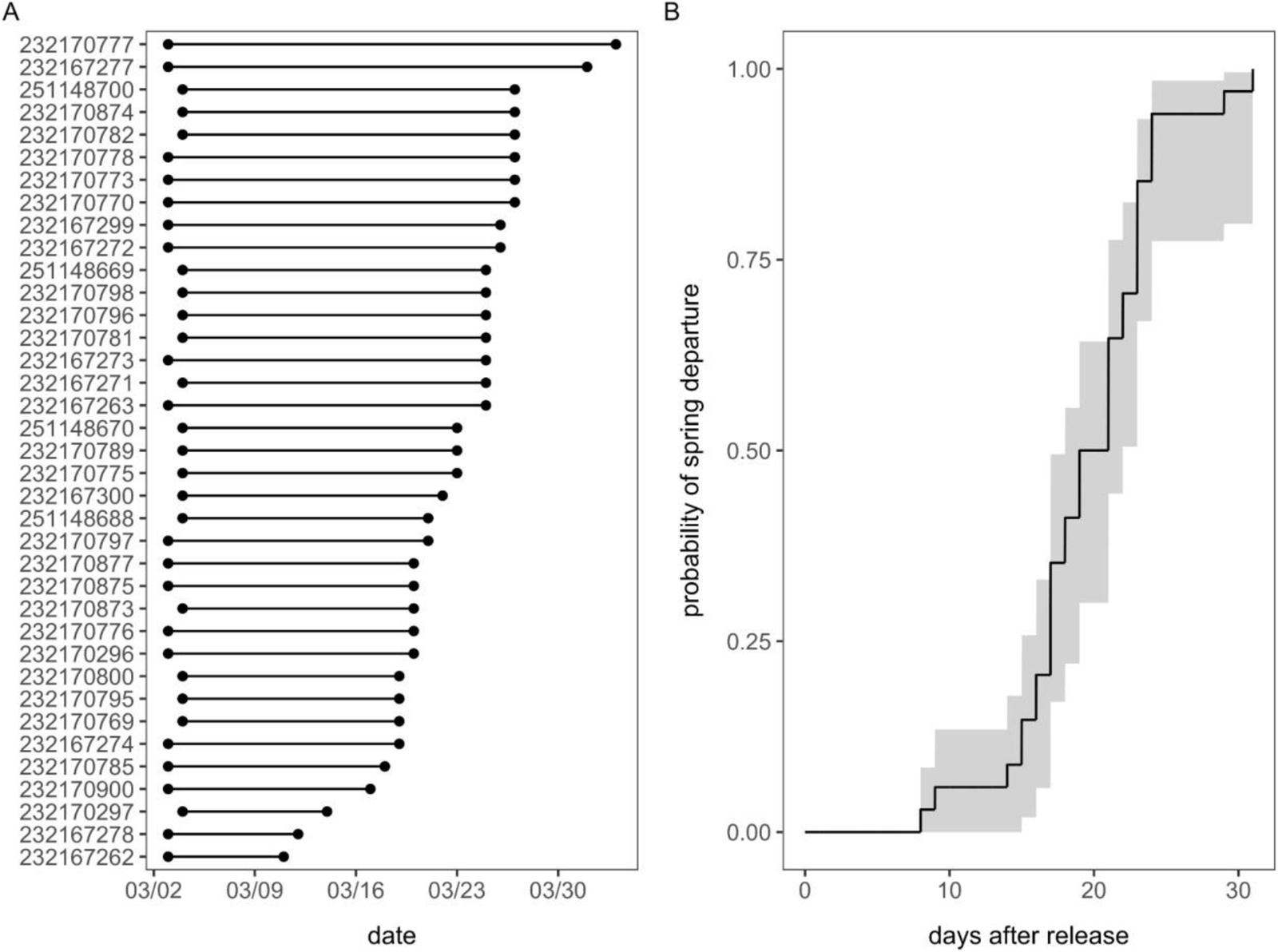
(A) Release and departure timing inferred from Motus at the Kent Farm Research Station for all 37 tagged juncos. (B) Mean probabilities of spring departure as inferred from Kaplan–Meier survival curves (i.e., 1 – survival probabilities) using the *survival* package alongside 95% confidence intervals.

**Table 1.**
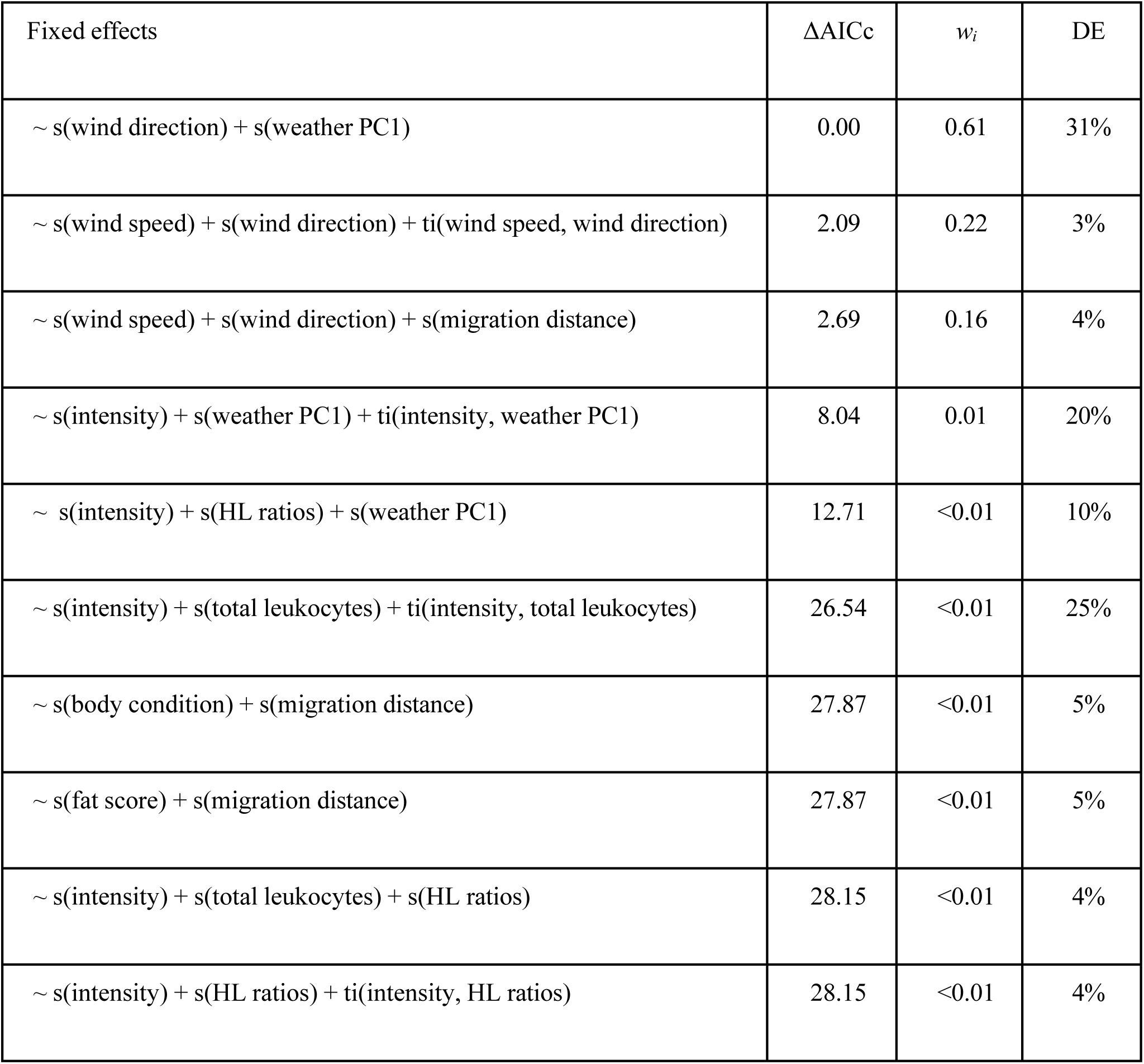
Comparison of CPHMMs predicting junco risk of spring departure; all models include a random effect of site. Candidate models are ranked by ΔAICc alongside Akaike weights (*w_i_*) and deviance explained (DE).

Our top model explained 30.5% of the deviance in spring migration risk, with both wind direction (χ21.8,2 = 17.42, p < 0.001) and weather PC1 (χ21.4,3 = 29.61, p < 0.001) being significant non-linear predictors of departure. Specifically, the risk of spring departure was greatest with average nightly southwestern winds and for humid and cool nights (Fig. 3).

**Figure 3.**
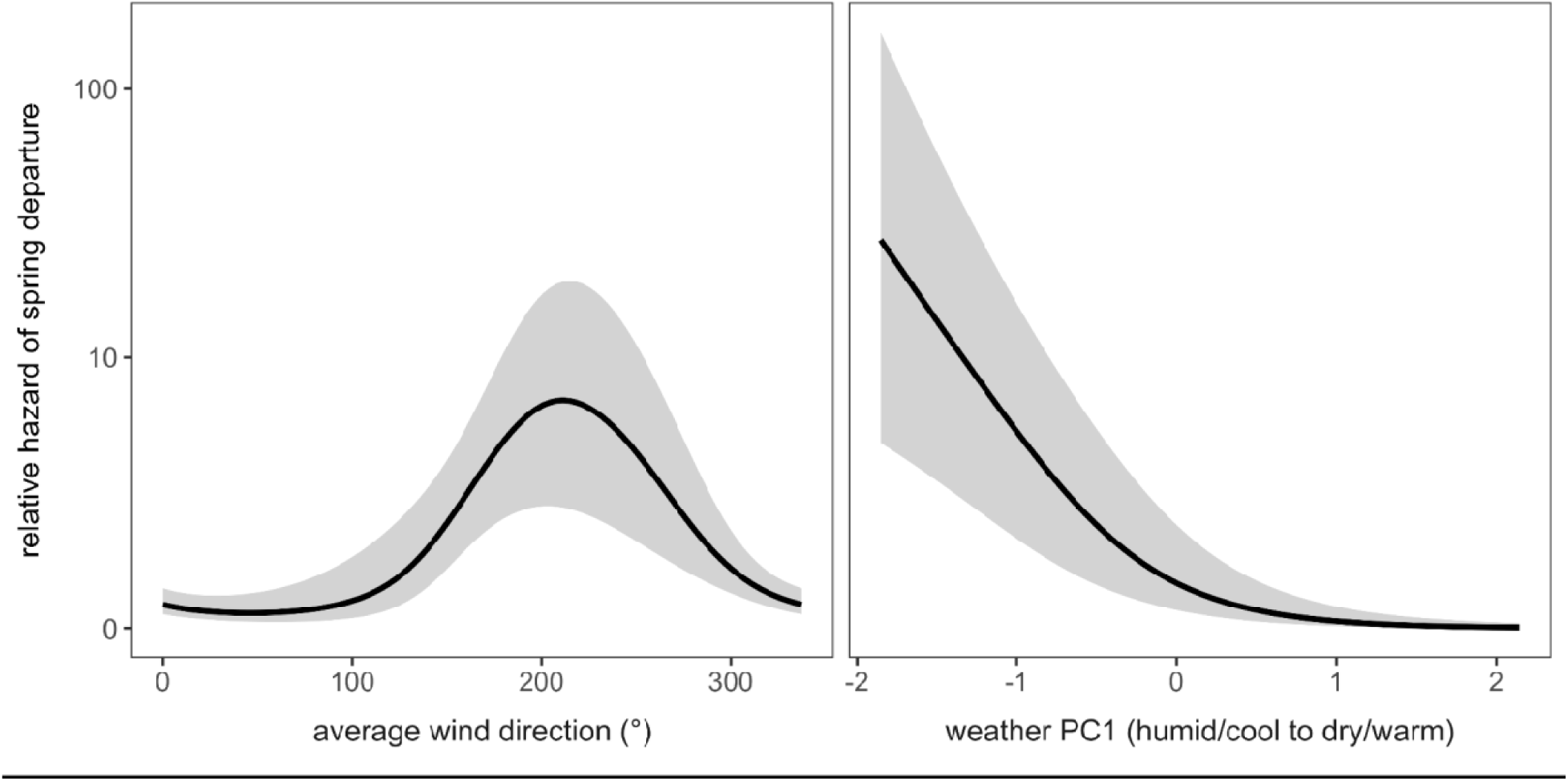
Relative hazards of spring departure estimated from the most competitive CPHMM as selected through AICc (*w_i_* = 0.61). The predicted relative hazard and 95% confidence intervals from this model are displayed for mean wind direction and weather PC1.

## Discussion

Numerous factors influence the timing of spring departure in birds, yet the relative influence of intrinsic and extrinsic variables on migration initiation remains a topic of debate. In this study, we tested the relative influence of these diverse factors on the risk of spring departure.in a modestly-sized group of wild-caught juncos. Weather variables outperformed all tested intrinsic variables. We predicted that both wind direction and speed would have the greatest influence on departure; however, only wind direction was supported by our top model. Temperature and relative humidity (i.e., weather PC1) also influenced departures, with humid and cool nights having greater risk of departure than warmer, drier nights. While we predicted that birds with intense haemosporidian infection would depart later than uninfected individuals or those with low-intensity infections, infection intensity did not influence departure risk. Additionally, higher fat reserves, lower leukocyte counts, and lower HL ratios did not affect departure risk. Our results suggest that the advantage of following winds outweigh the cost of haemosporidian infections and the other measured physiological variables during spring migration.

Many migratory birds harbor haemosporidian infections throughout the year, including during migration, (Ricklefs et al. 2017; Cornelius et al. 2014; Pulgarín-R et al. 2019) and ongoing research continues to clarify the effects of avian malaria on migration timing, duration, and distance. Prior work on juncos found greater prevalence of haemosporidian infections in a non-migratory subspecies compared to sympatric overwintering migrants following fall migration, suggesting that infected birds may be more likely to experience disease-induced mortality during migration (Slowinski et al., 2018). Similarly, juncos with longer migrations were more likely to show elevated HL ratios upon arrival to their wintering grounds; this effect was stronger in haemosporidian-infected birds, suggesting interactive effects of migration and infection on host energetics (Becker et al., 2019). Therefore, we predicted that juncos with more intense infections might delay migration initiation to mitigate these potential physiological costs. However, our results indicate that weather has a strong impact on migration irrespective of haemosporidian intensity. Although birds must navigate both intrinsic and extrinsic factors when choosing when to depart for migration, the cost of initiating spring migration under sub-optimal wind conditions may be greater than the cost of departing with a more intense haemosporidian infection. Importantly, our study focused on spring migration in adult juncos. While chronic haemosporidian infections may not impact migration timing during this life stage, it is unclear if this pattern would hold true for juncos during autumn migration, especially juveniles. In fact, juvenile European robins (Erithacus rubecula) with haemosporidian infections arrive at wintering ground later than uninfected subadults (Ágh et al. 2019). Thus, season, age, and stage of infection may influence whether haemosporidian infections delay migration initiation. Experimental infections in conjunction with departure tracking would provide more conclusive data on how weather interacts with parasite intensity to influence migration initiation.

Our analysis also indicates that migration distance and mean wind speed were relatively less informative predictors of spring departure date. Regarding migration distance, we expected that birds with longer spring migrations would depart earlier, as these individuals would potentially need to undertake prolonged stopovers to rest and refuel on the way to their breeding grounds. However, research suggests that long-distance migrants have a greater ability to modify migratory behavior while en route and thus departure date may not be directly related to migration distance as migration speed and routes are flexible (La Sorte and Fink, 2017; Marra et al., 2005). While juncos have been well-studied in regard to differential migration and migration timing (Cristol et al., 1999; Ketterson and Nolan, 1976), more research is needed to understand how distance to breeding ground influences departure date. Isotopic feather analysis is advancing work in this field, as is the improvement and proliferation of nanotag technology (e.g., Motus, Cellular Tracking Technologies), which will help further distinguish the various influences on migration distance as it relates to departure.

Past studies have shown that both wind speed and direction are important predictors of departure and that tail winds can significantly increase flight times and greatly decrease energy requirements (T. Alerstam, 1979; Bloch and Bruderer, 1982; Kemp et al., 2010; Liechti, 2006). There are several possibilities as to why these variables were less important in our analysis. Optimal wind selectivity is a product of many factors, including the size of the bird; rate of fat use; distance, duration, and altitude of migration; and even the wind pattern itself (Thomas Alerstam, 1979). Lastly, birds here departed over a relatively short time frame (a minimum of eight days after release, up to 31 days); therefore, there may not have been sufficient variation in wind speed and direction to outrank other predictors in our models.

In contrast to our expectations about physiological state and departure date, body condition, fat score, HL ratios, and leukocyte counts were also all uninformative predictors of spring migration date. Higher fat reserves are instrumental to migratory performance in songbirds (Price 2010) and may promote the onset of migratory restlessness, advancing their departure (Lupi et al., 2017; Studds and Marra, 2005). The lack of a relationship between physiological measures and departure date observed here could reflect individual variation in migration strategies; for example, strong selective pressure to arrive early at the breeding grounds may motivate some birds to leave in relatively poor body condition and with high HL ratios and then compensate during stopover (Prop et al., 2003). Lastly, the time between release from captivity and the onset of spring migration (8-31 days) may have precluded a true representation of some physiological measures at departure; for example, maximum fat deposition rates in migratory songbirds have been reported as high as 12% differences in lean body mass in one day (Lindström, 1991).

The findings presented here add to the large body of information on the importance of weather on migration departure in songbirds. Importantly, our study focused on migratory departure in male juncos, while females may be differentially impacted by the variables measured. Indeed, this species shows sex-based variation in overwintering latitudes, with females migrating earlier and further south than males (Nolan and Ketterson 1990). Our sample size is a significant limitation in understanding and asking these questions, as well. A more robust sample size would have increased the confidence in our findings. Additional data are also needed to understand whether the results of our study are context dependent; for example, intrinsic factors might become important during years with low food availability or inclement weather. This will be increasingly important for wildlife conservation going forward, as climate change is associated with increasingly severe temperatures and unpredictable weather events (X. Zhang et al. 2013; Stott 2016).

## Supporting information

Supp. Appx A, Table S1

## Data availability

The data that support the findings of this study are openly available in Dryad (https://datadryad.org) at https://datadryad.org/stash/share/Ne24zAcw7YGshc_QeUDPeWFrXNzJaPPGd-0kctpgK7Y.

## Declarations

Any use of trade, firm, or product names is for descriptive purposes only and does not imply endorsement by the U.S. Government.

## Ethics

Trapping, banding, and transmitter attachment was overseen by AJB and performed by AJB, DJB, and KMT under United States Federal Permit # 20261, Indiana state permit # 20-528, and approved under Bloomington Institutional Animal Care and Use Committee (BIACUC) # 18-030-15, Section 1. A Doctor of Veterinary Medicine (DVM) from Indiana University’s Laboratory Animal Resources (LAR) taught the oral gavage technique to AJB, DJB and KMT and this procedure was approved under the same protocol (BIACUC #18-030-15, Section 30).

## Acknowledgements

We thank Benjamin Higgins, Indiana University for his dedication in the lab. We also thank Peter Sauer and Kathyrn Evans in the Stable Isotope Research Facility at Indiana University for assistance in isotopic preparation and analysis. Lastly, we thank members of the Ketterson laboratory at Indiana University and two anonymous reviewers for helpful feedback on previous versions of this manuscript.

