## Supplementary material for "Determinants of spring migration departure dates in a New World sparrow: weather variables reign supreme": Supp. Appx A, Table S1

2 **reign supreme**
3 Allison J. Byrd^1,2*^, Katherine M. Talbott^2^, Tara M. Smiley^3^, Taylor B. Verrett^5^, Michael S. Gross^4^,
4 Michelle L. Hladik^4^, Ellen D. Ketterson^1,2^, Daniel J. Becker^5*^
5 ^1^Environmental Resilience Institute, Indiana University, Bloomington, IN
6 ^2^Department of Biology, Indiana University, Bloomington, IN
7 ^3^Department of Ecology and Evolution, Stony Brook University, Stony Brook, NY
8 ^4^U.S. Geological Survey, California Water Science Center, Sacramento, CA, 95819
9 ^5^School of Biological Sciences, University of Oklahoma, Norman, OK
10 *These authors contributed equally.
12 Building, 1001 E 3rd St, Bloomington, IN 47405

### 13 Funding

14 Funds were provided by Indiana University’s Grand Challenge Initiative, Prepared for
15 Environmental Change to EDK and TMS and from the University of Oklahoma to DJB.

#### 16 Conflict of interest

17 All authors declare they have no conflicts of interest.
18 **Keywords:** movement ecology, songbirds, haemosporidians, Motus, stable isotopes

#### 19 Abstract

20 Numerous factors influence the timing of spring migration in birds, yet the relative importance of intrinsic
21 and extrinsic variables on migration initiation remains unclear. To test for interactions among weather,
22 migration distance, parasitism, and physiology in determining spring departure date, we used Dark-eyed
23 Juncos (*Junco hyemalis hyemalis*) as a model migratory species known to harbor diverse and common
24 haemosporidian parasites. Prior to spring migration departure from their wintering grounds in Indiana,
25 USA, we quantified the intrinsic variables of fat, body condition (i.e., mass~tarsus residuals),
26 physiological stress (i.e., ratio of heterophils to lymphocytes), cellular immunity (i.e., leukocyte
27 composition and total count), migration distance (i.e., distance to the breeding grounds) using stable
28 isotopes of hydrogen from feathers, and haemosporidian parasite intensity. We then attached nanotags to
29 determine the timing of spring migration departure date using the Motus Wildlife Tracking System. We
30 used additive Cox proportional hazard mixed models to test how risk of spring migratory departure was
31 predicted by the combined intrinsic measures, along with meteorological predictors on the evening of
32 departure (i.e., average wind speed and direction, relative humidity, and temperature). Model comparisons
33 found that the best predictor of spring departure date was average nightly wind direction and a principal
34 component combining relative humidity and temperature. Juncos were more likely to depart for spring
35 migration on nights with largely southwestern winds and on warmer and drier evenings (relative to cooler
36 and more humid evenings). Our results indicate that weather conditions at take-off are more critical to
37 departure decisions than the measured physiological and parasitism variables.

38

39

#### 40 Introduction

41 Over-wintering duration and timing of departure for avian spring migration is a process influenced by a
42 host of extrinsic and intrinsic factors (Covino et al., 2015; Hurlbert and Liang, 2012; Satterfield et al.,
43 2018). The timing of spring migration must balance the advantages of early arrival on the breeding
44 grounds, including higher quality mates and territories (Gunnarsson et al., 2006; Møller, 1994; Newton,
45 2010; Rotics et al., 2018; Smith and Moore, 2005), with the risk of arriving before adequate food
46 resources are available (Gunnarsson et al., 2006; Møller, 1994; Newton, 2010; Rotics et al., 2018; Smith
47 and Moore, 2005). Stressors experienced on the wintering grounds can also affect the timing and duration
48 of spring migration. Extrinsic factors such as changing weather conditions (Marra et al., 2005) and
49 intrinsic factors such as high parasite intensity (Dietsch, 2005; Reed et al., 2003) can alter stopover
50 duration as birds recover from or accommodate these additional physiological burdens (Schmaljohann et
51 al. 2022; Skrip et al. 2015; Linscott and Senner 2021). The relative contribution of extrinsic and intrinsic
52 factors experienced at the wintering grounds in shaping spring departure remains less well understood,
53 and studies addressing these interactions are especially needed as anthropogenic pressures continue to
54 influence the climate and natural habitat (Kubelka et al., 2022; Visser et al., 2009; Wilcove and Wikelski,
55 2008).
56 For decades, ornithologists have known that weather variables influence avian migration timing in both
57 spring and fall (see:(Richardson, 1978). Factors that influence the probability of migratory departure
58 include wind speed (Chapman et al., 2016; Drake et al., 2014; Kemp et al., 2010; Liechti, 2006;
59 Nussbaumer et al., 2022), wind direction (Covino et al., 2015; Hebrard, 1971; Horton et al., 2016; Kemp
60 et al., 2010; Lack, 1963; Lack and Eastwood, 1962; Sinelschikova et al., 2007), temperature (Hüppop and
61 Winkel, 2006; Marra et al., 2005; Saino et al., 2007; Tøttrup et al., 2010; Usui et al., 2017), and relative
62 humidity (Klaassen et al., 2012; Liechti, 2006; Schmaljohann et al., 2009; Serra-Cobo et al., 1998; Zhang
63 and Wu, 2018). The high correlation among these weather variables adds to the difficulty in disentangling
64 the influence of these factors from potential additional stressors. For example, warm air can hold more
65 moisture than cold air, thus at the same absolute humidity, cooler air (perhaps counterintuitively) has
66 higher relative humidity than warmer air (Lawrence, 2005).
67 The relationship between wind and bird migration has been studies observed for decades, and prevailing
68 winds were a likely a selection factor in that shaping the evolution of migratory patterns (Alerstam 1979;
69 Alerstam 1979; Evans 2009; Able 1972; Richardson 1978). Birds that are highly selective of favorable
70 winds can maximize flight speed and minimize energy expenditure across both long and short-distance
71 migratory flights (Alerstam 1979). Migrants have faster ground and air-speeds in spring migration than
72 fall (Horton et al. 2016), and spring migration is typically completed over a shorter duration than fall
73 (Newton 2010; La Sorte et al. 2013; La Sorte et al. 2016; Nilsson et al. 2013). As such, models that
74 forecast bird migration rely heavily on wind data (Erni et al., 2002; Van Doren and Horton, 2018).
75 Intrinsic factors have also been recognized to affect readiness for migratory departure. Given the
76 energetic costs of migration (Wikelski et al., 2003), adequate fat stores are integral in providing fuel for
77 long-distance flight and larger stores can be a reliable predictor of migratory readiness(Price, 2010;
78 Ramenofsky, 1990; Weber et al., 1994; Witter and Cuthill, 1993); correspondingly, measures of body
79 condition (i.e., mass~tarsus residuals) allow for standardized body size comparisons among individuals
80 (Labocha and Hayes, 2012). The degree of physiological stress and immune state of individuals at the
81 wintering grounds can also affect spring migratory timing. Preparation for long-distance migration
82 increases plasma concentrations of corticosterone (the primary avian glucocorticoid responsible for
83 maintaining homeostasis) in many songbirds (Holberton, 1999; Landys et al., 2004), which can facilitate
84 earlier departure from the wintering grounds or spring stopover (Eikenaar et al., 2013, 2017). Elevated
85 corticosterone can shift the composition of leukocytes in blood, subsequently elevating the ratio of
86 heterophils to lymphocytes (HL ratios; Davis et al., 2008). While plasma corticosterone levels are highly
87 sensitive to the acute stress of capture, stress-induced changes in leukocyte profiles occur more slowly,
88 such that HL ratios can serve as a more tractable approximation of energetic costs prior to migration
89 (Davis and Maney, 2018). In a similar fashion, the energetic cost of migratory preparation can induce
90 trade-offs with the immune system (Lochmiller and Deerenberg, 2000), such that migrants may
91 downregulate immune activity or rely primarily on less costly immune defenses. For example, several
92 thrush species show lower total leukocyte counts upon arrival at stopover sites in spring, reflecting such
93 trade-offs (Owen and Moore, 2008). Variation in immune investment may in turn affect departure
94 decisions; recent work demonstrated that songbirds with higher titers of natural antibodies and
95 immunoglobulin Y have longer spring stopovers (Brust et al., 2022).
96 Related to seasonal variation in physiological stress and immunity, songbirds may also experience
97 additional overwintering stressors that affect spring migration. For example, exposure to toxins such as
98 neonicotinoids in wintering food sources (Eng et al., 2017; Goulson, 2014) can delay migration departure
99 or increase stopover duration as birds recover from or accommodate these burdens. An additional such
100 stressor could be through parasite infection, such as that from dipteran-borne haemosporidian blood
101 parasites (i.e., the genera *Plasmodium*, *Haemoproteus*, and *Leucocytozoon*). Experimental infections of
102 captive birds have shown migration-relevant physiological costs of infection, such as reduced general
103 activity (Mukhin et al., 2016; Yorinks and Atkinson, 2000) and body condition (Atkinson et al., 1995) as
104 well as shifts in the development of migratory restlessness (Kelly et al., 2016, 2020). Observational
105 studies have also identified impacts of haemosporidian infection on body condition in migrating birds
106 (Garvin et al., 2006; Merrill et al., 2018), on the timing of arrival to the breeding grounds (Asghar et al.,
107 2011; Santiago-Alarcon et al., 2013), and on autumn departure timing (Ágh et al., 2019). However, it
108 remains unclear how haemosporidian infections interact with extrinsic and other intrinsic factors to affect
109 migration timing.
110 In this study, we used overwintering migratory Dark-eyed Juncos (*Junco hyemalis hyemalis;* hereafter
111 “junco”) to test the hypothesis that physiological state and haemosporidian infection interact with weather
112 conditions and migration distance to shape spring migratory departure timing. Haemosporidia have been
113 well characterized in Dark-eyed juncos, with chronic (i.e., long- term) infections detected in wintering
114 and breeding populations (Becker et al. 2020; Slowinski et al. 2018; Talbott et al. 2022; Martínez-Renau
115 et al. 2022; Becker et al. 2019; Deviche, Greiner, and Manteca 2001; Ferrer 2022). We used stable
116 isotopes of hydrogen from feathers to estimate the distance to breeding grounds for each individual
117 (Bowen et al., 2014; Hobson, 1999; Hobson et al., 2012; Rubenstein and Hobson, 2004; Wunder, 2010)
118 and the Motus Wildlife Tracking System to determine departure date (Taylor et al., 2017). We predicted
119 that birds with higher fat reserves, low intensity or no haemosporidian parasitism, lower HL ratios, and
120 lower total leukocyte counts would depart earlier in the year and during evenings of following winds (i.e.,
121 tailwind). We also predicted that wind direction (following winds) would be more important than wind
122 speed, and that headwinds would be the strongest deterrent to departure, even more so than intrinsic
123 predictors.

#### 124 Methods

125 *Junco sampling and captive housing*
126 We caught wild juncos from 31 January to 26 February 2020 at four locations within 15 km of
127 Bloomington, Indiana. None of the birds were captured at the release site, and we systematically rotated
128 capture efforts through all four sites to avoid confounding results by capture date or location. We captured
129 juncos using baited mist nets and walk-in traps, took standard morphometric measurements including
130 mass and body fat (Fudickar et al., 2016; Jawor et al., 2006; Singh et al., 2019), and fitted each bird with
131 an aluminum US Fish & Wildlife Service leg band (federal permit # 20261, Indiana state permit 20-528).
132 While capture efforts were ongoing, juncos were held at an indoor aviary at Indiana University to prevent
133 additional exposure to arthropod vectors of haemosporidia. Juncos were provided *ad libitum* water and
134 food (mealworms, organic millet, organic sunflower seeds, and a blended mixture of organic millet,
135 organic carrots, and organic blueberries) and could fly freely in 6.4 x 3.2 m rooms. Light cycles for rooms
136 reflected the natural photoperiod of Bloomington, Indiana at the time of the study.
137 As a potential additional extrinsic factor, juncos in this study were randomly dosed with either a
138 neonicotinoid (imidacloprid suspended in sunflower seed oil) or control (sunflower seed oil) to
139 experimentally test impacts on migration timing. However, imidacloprid metabolites of 5-OH-imidacloprid
140 and imidacloprid-olefin were detected in only one bird’s post-dose plasma at 46.67 ng/mL and 72.23 ng/mL.
141 This individual was removed from all subsequent analyses. Imidacloprid or metabolites were not detected in
142 pre-dose or post-dose (6 hours after) plasma of any other individuals (*n* = 37; Supp. Appx A, Table S1).
143 Therefore, we did not include imidacloprid exposure as a possible predictor of migratory timing. Possible
144 explanations for the absence of imidacloprid detection in plasma samples are that the dose was
145 regurgitated or rapidly metabolized by all birds.
146 On the days of release (3 and 4 March 2020), we attached a nanotag to each bird (Lotek model NTQB2-2,
147 11 x 5 x 4 mm, 0.32g, ≦ 0.5 g total mass including leg harness). Birds were released at Kent Farm
148 Research Station (located nine miles east of Bloomington, IN), an area frequented by overwintering
149 juncos; we provided organic seed in feeders and on the ground in an attempt to minimize departure from
150 the area due to resource limitation (Bridge et al., 2010) .
151 *Hematological analyses*
152 At capture, we collected ≤ 150 μL blood from each bird by pricking the brachial vein with a sterile
153 needle, followed by collection with heparinized capillary tubes. We also collected the first secondary
154 feather from each bird for stable isotope analysis (Fudickar et al., 2016). We separated plasma and red
155 blood cells (centrifuged for 10 min at 10,000 rpm) and stored samples at -20℃ until DNA extraction
156 using a Maxwell RSC Whole Blood DNA Kit (Promega). Any birds unable to be sexed by wing length
157 and plumage were verified using sexing PCR of extracted DNA (Griffiths et al., 1998). Only males
158 (*n*=37) were included to control for sex differences in migration timing, as female juncos migrate earlier
159 than males (Nolan and Ketterson, 1990).
160 Because birds were held in captivity for variable lengths between capture and release with nanotags
161 (x̄ =25, range of 13 to 33 days), we collected additional blood samples 24 hours prior to release, using the
162 blood collection protocol described above. We prepared thin blood smears on glass slides stained with
163 Wright–Giemsa (Quick III, Astral Diagnostics). We then evaluated leucocyte profiles and
164 haemosporidian intensity using light microscopy (AmScope, B120C-E1). A single observer (TBV)
165 recorded the number of leukocytes under 400X magnification across 10 random fields to estimate
166 inflammatory state and investment in cellular immunity. A differential count was then performed by
167 recording the identity of the first 100 leukocytes (heterophils, lymphocytes, monocytes, eosinophils, and
168 basophils) at 1000X magnification (oil immersion;(Campbell, 1995). We then screened 100 fields of view
169 at 1000X magnification for *Plasmodium*, *Haemoproteus*, and *Leucocytozoon* *(Becker et al., 2019, 2020;*
170 *Cosgrove, 2005; Valkiunas et al., 2008)*. We derived the mean total leukocyte count, HL ratios, and the
171 total number of haemosporidian-infected erythrocytes (parasite intensity) as three predictor variables.
172 Usable blood smears were available for 34 of our 37 tagged individuals.
173 *Hydrogen isotope analysis*
174 We used stable isotopic analysis of hydrogen (δ^2^H) in feathers to infer probable breeding latitude and
175 median distance from capture point(Bowen et al., 2014; Hobson, 1999; Hobson et al., 2012; Rubenstein
176 and Hobson, 2004; Wunder, 2010). Feathers were cleaned using a 2:1 chloroform:methanol solution to
177 remove external oils and contaminants. We used a forced isotopic equilibration procedure to ensure
178 exchangeable hydrogen with a water vapor of known isotopic composition in a flow-through chamber
179 system at 115°C (Sauer et al., 2009; Schimmelmann, 1991). Samples were analyzed using a thermal
180 conversion element analyzer coupled with a ThermoFinnigan Delta Plus XP isotope ratio mass
181 spectrometer at the Indiana University Stable Isotope Research Facility. Isotopic data are reported in
182 standard per mil notation (‰) relative to VSMOW (Vienna Standard Mean Oceanic Water) using two
183 reference materials: USGS77 (polyethylene powder) and hexatriacontane 2 (C_36_ n-alkane 2). Analytical
184 precision was ± 1.0‰ for δ^2^H values. We calculated the isotopic composition of the non-exchangeable
185 hydrogen per sample, assuming a 17% exchangeability rate for feathers (Schimmelmann, 1991;
186 Schimmelmann et al., 1999).
187 We performed Bayesian geographic assignments for the breeding location of individual birds based on
188 feather δ^2^H values using the *assignR* package (Ma et al., 2020). To calibrate our precipitation-feather
189 isoscape, we used growing season precipitation isoscape rasters from waterisotopes.org (Bowen et al.,
190 2005; Bowen and Revenaugh, 2003) and the isotopic composition of non-migratory Dark-eyed juncos
191 from previous studies(Becker et al., 2019; Hobson et al., 2012). Median latitudinal and distance estimates
192 were calculated from geographic assignment probability maps by extracting the geographic cell
193 coordinates with the highest posterior probability (top 10%) using the *raster* package (Hijmans, 2019).
194 Migration distance was defined as the distance from the release site to the centroid of the estimated
195 breeding polygon (Becker et al., 2022; Wanamaker et al., 2020).
196 *Motus data processing*
197 Motus data (project 240; available at <https://motus.org/data>) were downloaded on 25 July 2020, four
198 months after the known period of spring migration in Indiana of wintering juncos (Ketterson and Nolan,
199 1982). Data were filtered and cleaned following a standard protocol using the *motus* package in R(Taylor
200 et al., 2017). Specifically, we removed runs with a low probability of being a true detection (i.e.,
201 motusFilter=0) and adopted a moderately strict filter to be more conservative about detection inclusions.
202 We then derived the departure date as the last day (ordinal date) a bird was detected at the Kent Farm
203 Research Station Motus station. We additionally report cases in which birds were detected at other Motus
204 stations between banding and our data download date.
205 *Weather data*
206 We obtained relative humidity data from ncei.noaa.gov (Diamond et al., 2013) and all remaining weather
207 variables (humidity, wind speed, wind direction [converted to wind rose] and weather type) from
208 timeanddate.com (Thorsen, 1995-2023). We averaged hourly data from 1800-2400 on departure date
209 evenings (thus, variables are labeled average humidity and so on).
210 Because average relative humidity and average temperature are dependent (Lawrence, 2005), we
211 conducted a principal components analysis (PCA) of these two variables, with variables centered and
212 scaled to have unit variance. The first PC (hereafter weather PC1) explained 56% of the variation and was
213 loaded negatively by average relative humidity (–0.71) and positively by average temperature (0.71);
214 resulting values indicate increasingly warm and dry evening weather. This variable was used in
215 downstream analyses alongside average wind speed and average wind direction.
216 *Statistical analyses*
217 Prior to analyses of migration timing, we used a linear mixed effects model with a random effect of site to
218 derive an index of body condition through the residuals of a regression of mass (measured prior to
219 release) on tarsus length (Schulte-Hostedde et al., 2005; Wanamaker et al., 2020).
220 We modeled days until spring migratory departure under an event-time analysis framework, using
221 additive Cox proportional hazard mixed models (CPHMMs, (Therneau and Grambsch, 2000) and
222 Grambsch, 2000). These semi-parametric models allow determining how the risk of departure changes
223 with covariates that can also vary with time, and such models have been applied to study migration timing
224 in both avian and non-avian systems (Castro-Santos and Haro, 2003; Dossman et al., 2015). We used the
225 *mgcv* package to fit CPHMMs with a random effect of site. To test the direct and interactive effects of
226 extrinsic (i.e., weather PC1, wind direction, wind speed) and intrinsic predictors (i.e., fat, body condition,
227 total leukocytes, HL ratios, haemosporidian intensity) on risk of spring departure (Wood, 2017).
228 Including all predictor variables and biologically relevant interactions in a single full model was not
229 possible given our sample size (*n* = 34, excluding birds without hematology data). We instead built 10
230 candidate CPHMMs representing *a priori* hypotheses of additive and interactive effects while restricting
231 models to at most three fixed effects to limit overfitting (Burnham and Anderson, 2002). Most predictors
232 showed low collinearity (ρ ranged from –0.63 to 0.62, x̄ = 0.01), and moderately correlated predictors
233 (i.e., weather PC1 and wind speed, ρ = –0.63; body fat and condition, ρ = 0.62) were excluded from the
234 same models. All predictors were modeled using thin plate splines with smoothing penalty with the
235 exception of wind direction, which used a cyclic cubic spline to account for circular data. Interaction
236 terms were modeled as tensor products. Our models represented additive effects of intrinsic variables only
237 (e.g., fat, haemosporidian intensity, and total leukocytes), additive effects of extrinsic variables only (e.g.,
238 wind direction and weather PC1), additive effects of both intrinsic and extrinsic variables (e.g.,
239 haemosporidian intensity, HL ratios, and weather PC1), interactive effects of intrinsic variables (e.g.,
240 effects of haemosporidian intensity dependent on HL ratios), interactive effects of extrinsic variables
241 (e.g., effects of wind speed depend on wind direction), and interactive effects of intrinsic and extrinsic
242 variables (e.g., effects of weather PC1 depend on haemosporidian intensity). We compared CPHMMs fit
243 with maximum likelihood using Akaike information criterion adjusted for small sample size (AICc) with
244 the *MuMIn* package(Bartoń, 2013). We also derived Akaike weights (*w_i_*) to facilitate comparison and
245 considered models within two ΔAICc of the top model to be competitive. Competitive models were
246 refitted to the full dataset, with CPHMM predictions visualized as relative hazards with 95% confidence
247 intervals (Nakagawa and Schielzeth, 2013).

#### 248 Results

249 *Hydrogen isotopes and likely breeding origins*
250 Hydrogen isotopic composition of junco feathers ranged from -158.9 to -95.8%, reflecting median
251 breeding locations from 66 to 51°N, respectively (Fig. 1). Maximum and minimum latitudinal estimates
252 spanned from 70 to 36 °N. Estimated distances to breeding grounds accordingly ranged from
253 approximately 2086 to 3900 kilometers (x̄ =3297, SE=78.4). Incidentally, we detected six of our 37 tagged
254 juncos at other Motus towers following their departure from Indiana (Fig. 1). Five birds were detected at
255 the Lake Petite station in Wisconsin (42.5117°, -88.5488°) and one bird was detected at the Werden
256 station (42.7551°, -80.2724°) in Ontario, Canada.
257 *Haemosporidian infection*
258 We identified haemosporidian infections in 11 of 34 juncos prior to release (32.4%, 95% CI: 19.1–
259 49.2%). Only two individuals had detectable *Haemoproteus* spp. infection (intensity=16–23 infected
260 erythrocytes from 100 fields of view), and nine individuals had detectable *Leucocytozoon* spp. infection
261 (intensity=1–7 infected erythrocytes from 100 fields of view); no individuals harbored co-infecting
262 haemosporidian parasites. We also opportunistically detected *Trypanosoma* spp. in a single bird (1/34,
263 two parasites were detected from 100 fields of view), and this individual was not infected by either
264 *Haemoproteus* spp. nor *Leucocytozoon* spp. parasites.
265 *Extrinsic and intrinsic predictors of migration timing*
266 All birds departed 8–31 days after release (x̄ =19.11 ± 0.81 SE), between March 11 and April 3 2020 (Fig.
267 2). Comparison among 10 CPHMMs predicting risk of departure date as a function of extrinsic and
268 intrinsic factors identified only one competitive model, which included the nonlinear additive effects of
269 wind speed and weather PC1 (*w_i_*=0.61; Table 1). Wind direction had the greatest relative importance
270 among predictors (99%), followed by weather PC1 (62%); wind speed, the interaction between wind
271 speed and weather PC1, and estimated migration distance all had lesser importance (38%, 22%, and 16%,
272 respectively). All intrinsic predictors (i.e., fat, body condition, total leukocytes, HL ratios) and their
273 interactions were unimportant (≤ 1%).
274 Our top model explained 30.5% of the deviance in spring migration risk, with both wind direction
275 (χ21.8,2 = 17.42, p < 0.001) and weather PC1 (χ21.4,3 = 29.61, p < 0.001) being significant non-linear
276 predictors of departure. Specifically, the risk of spring departure was greatest with average nightly
277 southwestern winds and for humid and cool nights (Fig. 3).

278

#### 279 Discussion

280 Numerous factors influence the timing of spring departure in birds, yet the relative influence of
281 intrinsic and extrinsic variables on migration initiation remains a topic of debate. In this study, we
282 tested the relative influence of these diverse factors on the risk of spring departure.in a modestly-
283 sized group of wild-caught juncos. Weather variables outperformed all tested intrinsic variables.
284 We predicted that both wind direction and speed would have the greatest influence on departure;
285 however, only wind direction was supported by our top model. Temperature and relative humidity
286 (i.e., weather PC1) also influenced departures, with humid and cool nights having greater risk of
287 departure than warmer, drier nights. While we predicted that birds with intense haemosporidian
288 infection would depart later than uninfected individuals or those with low-intensity infections,
289 infection intensity did not influence departure risk. Additionally, higher fat reserves, lower
290 leukocyte counts, and lower HL ratios did not affect departure risk. Our results suggest that the
291 advantage of following winds outweigh the cost of haemosporidian infections and the other
292 measured physiological variables during spring migration.
293 Many migratory birds harbor haemosporidian infections throughout the year, including during migration,
294 (Ricklefs et al. 2017; Cornelius et al. 2014; Pulgarín-R et al. 2019) and ongoing research continues to
295 clarify the effects of avian malaria on migration timing, duration, and distance. Prior work on juncos
296 found greater prevalence of haemosporidian infections in a non-migratory subspecies compared to
297 sympatric overwintering migrants following fall migration, suggesting that infected birds may be more
298 likely to experience disease-induced mortality during migration (Slowinski et al., 2018). Similarly, juncos
299 with longer migrations were more likely to show elevated HL ratios upon arrival to their wintering
300 grounds; this effect was stronger in haemosporidian-infected birds, suggesting interactive effects of
301 migration and infection on host energetics (Becker et al., 2019). Therefore, we predicted that juncos with
302 more intense infections might delay migration initiation to mitigate these potential physiological costs.
303 However, our results indicate that weather has a strong impact on migration irrespective of
304 haemosporidian intensity. Although birds must navigate both intrinsic and extrinsic factors when
305 choosing when to depart for migration, the cost of initiating spring migration under sub-optimal wind
306 conditions may be greater than the cost of departing with a more intense haemosporidian infection.
307 Importantly, our study focused on spring migration in adult juncos. While chronic haemosporidian
308 infections may not impact migration timing during this life stage, it is unclear if this pattern would hold
309 true for juncos during autumn migration, especially juveniles. In fact, juvenile European robins (Erithacus
310 rubecula) with haemosporidian infections arrive at wintering ground later than uninfected subadults (Ágh
311 et al. 2019). Thus, season, age, and stage of infection may influence whether haemosporidian infections
312 delay migration initiation. Experimental infections in conjunction with departure tracking would provide
313 more conclusive data on how weather interacts with parasite intensity to influence migration initiation.
314 Our analysis also indicates that migration distance and mean wind speed were relatively less informative
315 predictors of spring departure date. Regarding migration distance, we expected that birds with longer
316 spring migrations would depart earlier, as these individuals would potentially need to undertake
317 prolonged stopovers to rest and refuel on the way to their breeding grounds. However, research suggests
318 that long-distance migrants have a greater ability to modify migratory behavior while en route and thus
319 departure date may not be directly related to migration distance as migration speed and routes are flexible
320 (La Sorte and Fink, 2017; Marra et al., 2005). While juncos have been well-studied in regard to
321 differential migration and migration timing (Cristol et al., 1999; Ketterson and Nolan, 1976), more
322 research is needed to understand how distance to breeding ground influences departure date. Isotopic
323 feather analysis is advancing work in this field, as is the improvement and proliferation of nanotag
324 technology (e.g., Motus, Cellular Tracking Technologies), which will help further distinguish the various
325 influences on migration distance as it relates to departure.
326 Past studies have shown that both wind speed and direction are important predictors of departure
327 and that tail winds can significantly increase flight times and greatly decrease energy requirements
328 (T. Alerstam, 1979; Bloch and Bruderer, 1982; Kemp et al., 2010; Liechti, 2006). There are several
329 possibilities as to why these variables were less important in our analysis. Optimal wind selectivity
330 is a product of many factors, including the size of the bird; rate of fat use; distance, duration, and
331 altitude of migration; and even the wind pattern itself (Thomas Alerstam, 1979). Lastly, birds here
332 departed over a relatively short time frame (a minimum of eight days after release, up to 31 days);
333 therefore, there may not have been sufficient variation in wind speed and direction to outrank other
334 predictors in our models.
335 In contrast to our expectations about physiological state and departure date, body condition, fat score, HL
336 ratios, and leukocyte counts were also all uninformative predictors of spring migration date. Higher fat
337 reserves are instrumental to migratory performance in songbirds (Price 2010) and may promote the onset
338 of migratory restlessness, advancing their departure (Lupi et al., 2017; Studds and Marra, 2005). The lack
339 of a relationship between physiological measures and departure date observed here could reflect
340 individual variation in migration strategies; for example, strong selective pressure to arrive early at the
341 breeding grounds may motivate some birds to leave in relatively poor body condition and with high HL
342 ratios and then compensate during stopover (Prop et al., 2003). Lastly, the time between release from
343 captivity and the onset of spring migration (8-31 days) may have precluded a true representation of some
344 physiological measures at departure; for example, maximum fat deposition rates in migratory songbirds
345 have been reported as high as 12% differences in lean body mass in one day (Lindström, 1991).
346 The findings presented here add to the large body of information on the importance of weather on
347 migration departure in songbirds. Importantly, our study focused on migratory departure in male juncos,
348 while females may be differentially impacted by the variables measured. Indeed, this species shows sex-
349 based variation in overwintering latitudes, with females migrating earlier and further south than males
350 (Nolan and Ketterson 1990). Our sample size is a significant limitation in understanding and asking these
351 questions, as well. A more robust sample size would have increased the confidence in our findings.
352 Additional data are also needed to understand whether the results of our study are context dependent; for
353 example, intrinsic factors might become important during years with low food availability or inclement
354 weather. This will be increasingly important for wildlife conservation going forward, as climate change is
355 associated with increasingly severe temperatures and unpredictable weather events (X. Zhang et al. 2013;
356 Stott 2016).

#### 357 Data availability

358 The data that support the findings of this study are openly available in Dryad (https://datadryad.org) at
359 [https://datadryad.org/stash/share/Ne24zAcw7YGshc_QeUDPeWFrXNzJaPPGd-0kctpgK7Y.](https://datadryad.org/stash/share/Ne24zAcw7YGshc_QeUDPeWFrXNzJaPPGd-0kctpgK7Y)

### 360 Declarations

### 362 Any use of trade, firm, or product names is for descriptive purposes only and does not imply

### 363 endorsement by the U.S. Government.

### 364 Ethics

365 Trapping, banding, and transmitter attachment was overseen by AJB and performed by AJB, DJB, and
366 KMT under United States Federal Permit # 20261, Indiana state permit # 20-528, and approved under
367 Bloomington Institutional Animal Care and Use Committee (BIACUC) # 18-030-15, Section 1. A Doctor
368 of Veterinary Medicine (DVM) from Indiana University’s Laboratory Animal Resources (LAR) taught
369 the oral gavage technique to AJB, DJB and KMT and this procedure was approved under the same
370 protocol (BIACUC #18-030-15, Section 30).

### 371 Competing interests

372 The authors declare they have no competing interests.

### 373 Acknowledgements

374 We thank Benjamin Higgins, Indiana University for his dedication in the lab. We also thank
375 Peter Sauer and Kathyrn Evans in the Stable Isotope Research Facility at Indiana University for
376 assistance in isotopic preparation and analysis. Lastly, we thank members of the Ketterson
377 laboratory at Indiana University and two anonymous reviewers for helpful feedback on previous
378 versions of this manuscript.

375

419	244.

420 Castro-Santos T and Haro A (2003) Quantifying migratory delay: a new application of survival analysis
421 methods. *Canadian journal of fisheries and aquatic sciences. Journal canadien des sciences*
422 *halieutiques et aquatiques* 60(8). Canadian Science Publishing: 986–996.
423 Chapman JW, Nilsson C, Lim KS, et al. (2016) Adaptive strategies in nocturnally migrating insects and
424 songbirds: contrasting responses to wind. *The Journal of animal ecology* 85(1). Wiley Online
425 Library: 115–124.
426 Cornelius EA, Davis AK and Altizer SA (2014) How important are hemoparasites to migratory
427 songbirds? Evaluating physiological measures and infection status in three neotropical migrants
428 during stopover. *Physiological and biochemical zoology: PBZ* 87(5). journals.uchicago.edu: 719–
429 728.
430 Cosgrove C (2005) Avian malaria parasites and other haemosporidia.—Gediminas valkiunas. 2004. CRC
431 press, Boca Raton, Florida, USA. 932 pp. ISBN 0-415-30097-5. US$170 (hardcover). *Systematic*
432 *biology* 54(5). Oxford University Press (OUP): 860–863.
433 Covino KM, Holberton RL and Morris SR (2015) Factors influencing migratory decisions made by
434 songbirds on spring stopover. *Journal of avian biology* 46(1): 73–80.
435 Cristol DA, Baker MB and Carbone C (1999) Differential Migration Revisited. In: Nolan V, Ketterson
436 ED, and Thompson CF (eds) *Current Ornithology*. Boston, MA: Springer US, pp. 33–88.
437 Davis AK and Maney DL (2018) The use of glucocorticoid hormones or leucocyte profiles to measure
438 stress in vertebrates: What’s the difference? *Methods in ecology and evolution / British Ecological*
439 *Society* 9(6). Wiley: 1556–1568.
440 Davis AK, Maney DL and Maerz JC (2008) The use of leukocyte profiles to measure stress in vertebrates:
441 a review for ecologists. *Functional ecology* 22(5). Wiley: 760–772.
442 de Angeli Dutra D, Fecchio A, Braga ÉM, et al. (2022) Migratory behaviour does not alter
443 cophylogenetic congruence between avian hosts and their haemosporidian parasites. *Parasitology*.
444 cambridge.org: 1–8.
445 Deviche P, Greiner EC and Manteca X (2001) Seasonal and age-related changes in blood parasite
446 prevalence in Dark-eyed Juncos (Junco hyemalis, Aves, Passeriformes). *The Journal of experimental*
447 *zoology* 289(7). Wiley: 456–466.
448 Diamond HJ, Karl TR, Palecki MA, et al. (2013) U.S. Climate Reference Network after One Decade of
449 Operations: Status and Assessment. *Bulletin of the American Meteorological Society* 94(4).
450 American Meteorological Society: 485–498.
451 Dietsch TV (2005) Seasonal variation of infestation by ectoparasitic chigger mite larvae (Acarina:
452 Trombiculidae) on resident and migratory birds in coffee agroecosystems of Chiapas, Mexico. *The*
453 *Journal of parasitology* 91(6): 1294–1303.
454 Dossman BC, Mitchell GW, Norris DR, et al. (2015) The effects of wind and fuel stores on stopover
455 departure behavior across a migratory barrier. *Behavioral ecology: official journal of the*
456 *International Society for Behavioral Ecology* 27(2). Oxford Academic: 567–574.
457 Drake A, Rock CA, Quinlan SP, et al. (2014) Wind speed during migration influences the survival, timing
458 of breeding, and productivity of a neotropical migrant, Setophaga petechia. *PloS one* 9(5).
459 journals.plos.org: e97152.
460 Eikenaar C, Fritzsch A and Bairlein F (2013) Corticosterone and migratory fueling in Northern wheatears
461 facing different barrier crossings. *General and comparative endocrinology* 186. Elsevier: 181–186.
462 Eikenaar C, Müller F and Leutgeb C (2017) Corticosterone and timing of migratory departure in a
463 songbird. *Royal Society B …*. royalsocietypublishing.org. Epub ahead of print 2017.
464 Eng ML, Stutchbury BJM and Morrissey CA (2017) Imidacloprid and chlorpyrifos insecticides impair
465 migratory ability in a seed-eating songbird. *Scientific reports* 7(1): 15176.
466 Ephrath JE, Goudriaan J and Marani A (1996) Modelling diurnal patterns of air temperature, radiation
467 wind speed and relative humidity by equations from daily characteristics. *Agricultural systems* 51(4).
468 Elsevier: 377–393.
469 Erni B, Liechti F, Underhill LG, et al. (2002) Wind and rain govern the intensity of nocturnal bird
470 migration in Central Europe- a log-linear regression analysis. *Ardea* 90(1). researchgate.net: 155–
471 166.
472 Ferrer MM (2022) *AVIAN HAEMOSPORIDIAN BLOOD PARASITE DIVERSITY, PREVALENCE, AND*
473 *DISTRIBUTION IN MICHIGAN’S WESTERN UPPER PENINSULA*. Michigan Technological
474 University. Available at: <https://digitalcommons.mtu.edu/etdr/1470/> (accessed 7 June 2023).
475 Fudickar AM, Greives TJ, Atwell JW, et al. (2016) Reproductive Allochrony in Seasonally Sympatric
476 Populations Maintained by Differential Response to Photoperiod: Implications for Population
477 Divergence and Response to Climate Change. *The American naturalist* 187(4): 436–446.
478 Garvin MC, Szell CC and Moore FR (2006) BLOOD PARASITES OF NEARCTIC–NEOTROPICAL
479 MIGRANT PASSERINE BIRDS DURING SPRING TRANS-GULF MIGRATION: IMPACT ON
480 HOST BODY CONDITION. *The Journal of parasitology* 92(5). Allen Press: 990–996.
481 Griffiths R, Double MC, Orr K, et al. (1998) A DNA test to sex most birds. *Molecular ecology* 7(8).
482 Wiley Online Library: 1071–1075.
483 Gunnarsson TG, Gill JA, Atkinson PW, et al. (2006) Population-scale drivers of individual arrival times
484 in migratory birds. *The Journal of animal ecology* 75(5): 1119–1127.
485 Hahn S, Bauer S, Dimitrov D, et al. (2018) Low intensity blood parasite infections do not reduce the
486 aerobic performance of migratory birds. *Proceedings. Biological sciences / The Royal Society*
487 285(1871). royalsocietypublishing.org.
488 Hallmann CA, Foppen RPB, van Turnhout CAM, et al. (2014) Declines in insectivorous birds are
489 associated with high neonicotinoid concentrations. *Nature* 511(7509): 341–343.
490 Hebrard JJ (1971) The nightly initiation of passerine migration in spring: a direct visual study. *The Ibis*.
491 Wiley Online Library. Epub ahead of print 1971. DOI: [10.1111/j.1474-919X.1971.tb05119.x](http://dx.doi.org/10.1111/j.1474-919X.1971.tb05119.x).
492 Hijmans RJ (2019) Introduction to the’raster’package (version 2.8-19). mran.microsoft.com. Epub ahead
493 of print 2019.
494 Hobson KA (1999) Tracing origins and migration of wildlife using stable isotopes: a review. *Oecologia*

495	120(3): 314–326.

496 Hobson KA, Van Wilgenburg SL, Wassenaar LI, et al. (2012) Linking hydrogen (δ2H) isotopes in
497 feathers and precipitation: sources of variance and consequences for assignment to isoscapes. *PloS*
498 *one* 7(4): e35137.
499 Holberton RL (1999) Changes in patterns of corticosterone secretion concurrent with migratory fattening
500 in a neotropical migratory bird. *General and comparative endocrinology* 116(1). Elsevier: 49–58.
501 Horton KG, Van Doren BM, Stepanian PM, et al. (2016) Seasonal differences in landbird migration
502 strategies. *The Auk* 133(4). Oxford Academic: 761–769.
503 Hüppop O and Winkel W (2006) Climate change and timing of spring migration in the long-distance
504 migrant Ficedula hypoleuca in central Europe: the role of spatially different temperature changes
505 along migration routes. *Journal of ornithology / DO-G* 147(2). Springer: 344–353.
506 Hurlbert AH and Liang Z (2012) Spatiotemporal variation in avian migration phenology: citizen science
507 reveals effects of climate change. *PloS one* 7(2): e31662.
508 Jawor JM, McGlothlin JW, Casto JM, et al. (2006) Seasonal and individual variation in response to
509 GnRH challenge in male dark-eyed juncos (Junco hyemalis). *General and comparative*
510 *endocrinology* 149(2). Elsevier: 182–189.
511 Kelly TR, MacGillivray HL, Sarquis-Adamson Y, et al. (2016) Seasonal migration distance varies with
512 natal dispersal and predicts parasitic infection in song sparrows. *Behavioral Ecology and*
513 *Sociobiology*. Available at: <http://dx.doi.org/10.1007/s00265-016-2191-2>.
514 Kelly TR, Rubin BD, MacDougall-Shackleton SA, et al. (2020) Experimental Malaria Infection Affects
515 Songbirds’ Nocturnal Migratory Activity. *Physiological and biochemical zoology: PBZ* 93(2). The
516 University of Chicago Press: 97–110.
517 Kemp MU, Shamoun-Baranes J, Van Gasteren H, et al. (2010) Can wind help explain seasonal
518 differences in avian migration speed? *Journal of avian biology* 41(6). Wiley: 672–677.
519 Ketterson ED and Nolan V (1982) The Role of Migration and Winter Mortality in the Life History of a
520 Temperate-Zone Migrant, the Dark-Eyed Junco, as Determined from Demographic Analyses of
521 Winter Populations. *The Auk* 99(2). Oxford Academic: 243–259.
522 Ketterson ED and Nolan V Jr (1976) Geographic variation and its climatic correlates in the sex ratio of
523 eastern-wintering dark-eyed juncos (junco hyemalis hyemalis). *Ecology* 57(4). Wiley: 679–693.
524 Klaassen M, Hoye BJ, Nolet BA, et al. (2012) Ecophysiology of avian migration in the face of current
525 global hazards. *Philosophical transactions of the Royal Society of London. Series B, Biological*
526 *sciences* 367(1596): 1719–1732.
527 Kubelka V, Sandercock BK, Székely T, et al. (2022) Animal migration to northern latitudes:
528 environmental changes and increasing threats. *Trends in ecology & evolution* 37(1). Elsevier: 30–41.
529 Labocha MK and Hayes JP (2012) Morphometric indices of body condition in birds: a review. *Journal of*
530 *ornithology / DO-G* 153(1). Springer: 1–22.
531 Lack D (1963) Weather factors initiating migration. *Proc. XIII Int. Ornithol. Congr*. Epub ahead of print

532	1963.

533 Lack D and Eastwood E (1962) Radar films of migration over eastern England. *British birds; an*
534 *illustrated magazine devoted to the birds on the British list*. Epub ahead of print 1962.
535 Landys MM, Wingfield JC and Ramenofsky M (2004) Plasma corticosterone increases during migratory
536 restlessness in the captive white-crowned sparrow Zonotrichia leucophrys gambelli. *Hormones and*
537 *behavior* 46(5). Elsevier: 574–581.
538 La Sorte FA and Fink D (2017) Migration distance, ecological barriers and en-route variation in the
539 migratory behaviour of terrestrial bird populations. *Global ecology and biogeography: a journal of*
540 *macroecology* 26(2). Wiley: 216–227.
541 Lawrence MG (2005) The Relationship between Relative Humidity and the Dewpoint Temperature in
542 Moist Air: A Simple Conversion and Applications. *Bulletin of the American Meteorological Society*
543 86(2). American Meteorological Society: 225–234.
544 Liechti F (2006) Birds: blowin’ by the wind? *Journal of ornithology / DO-G* 147(2). Springer Science and
545 Business Media LLC: 202–211.
546 Lindström Å (1991) Maximum Fat Deposition Rates in Migrating Birds. *Ornis Scandinavica* 22(1).
547 [[Nordic Society Oikos, Wiley]: 12–19.](http://paperpile.com/b/9rwraW/NbBgg)
548 Lochmiller RL and Deerenberg C (2000) Trade-offs in evolutionary immunology: just what is the cost of
549 immunity? *Oikos* 88(1). Wiley: 87–98.
550 Lopez-Antia A, Ortiz-Santaliestra ME, Mougeot F, et al. (2013) Experimental exposure of red-legged
551 partridges (Alectoris rufa) to seeds coated with imidacloprid, thiram and difenoconazole.
552 *Ecotoxicology* 22(1). Springer: 125–138.
553 Lopez-Antia A, Ortiz-Santaliestra ME, Mougeot F, et al. (2015) Imidacloprid-treated seed ingestion has
554 lethal effect on adult partridges and reduces both breeding investment and offspring immunity.
555 *Environmental research* 136. Elsevier: 97–107.
556 Lupi S, Maggini I, Goymann W, et al. (2017) Effects of body condition and food intake on stop-over
557 decisions in Garden Warblers and European Robins during spring migration. *Journal of ornithology*
558 */ DO-G* 158(4). Springer: 989–999.
559 Ma C, Vander Zanden HB, Wunder MB, et al. (2020) assignR : An r package for isotope‐based
560 geographic assignment. *Methods in ecology and evolution / British Ecological Society* 11(8). Wiley:
561 996–1001.
562 Marra PP, Francis CM, Mulvihill RS, et al. (2005) The influence of climate on the timing and rate of
563 spring bird migration. *Oecologia* 142(2): 307–315.
564 Martínez-Renau E, Rojas-Estévez N, Friis G, et al. (2022) Haemosporidian parasite diversity and
565 prevalence in the songbird genus Junco across Central and North America. *Ornithology* 139(3).
566 Oxford Academic: ukac022.
567 Mason R, Tennekes H, Sánchez-Bayo F, et al. (2013) Immune suppression by neonicotinoid insecticides
568 at the root of global wildlife declines. *J Environ Immunol Toxicol* 1(1): 3–12.
569 Merrill L, Levengood JM, England JC, et al. (2018) Blood parasite infection linked to condition of
570 spring-migrating Lesser Scaup (Aythya affinis). *Canadian journal of zoology* 96(10). Canadian
571 Science Publishing: 1145–1152.
572 Møller AP (1994) Phenotype-dependent arrival time and its consequences in a migratory bird. *Behavioral*
573 *ecology and sociobiology* 35(2). Springer Nature: 115–122.
574 Mukhin A, Palinauskas V, Platonova E, et al. (2016) The Strategy to Survive Primary Malaria Infection:
575 An Experimental Study on Behavioural Changes in Parasitized Birds. *PloS one* 11(7).
576 journals.plos.org: e0159216.
577 Nakagawa S and Schielzeth H (2013) A general and simple method for obtaining R2 from generalized
578 linear mixed-effects models. *Methods in ecology and evolution / British Ecological Society* 4(2).
579 Wiley Online Library: 133–142.
580 Newton I (2010) *The Migration Ecology of Birds*. Elsevier.
581 Nolan V Jr and Ketterson ED (1990) Timing of autumn migration and its relation to winter distribution in
582 dark-eyed juncos. *Ecology* 71(4). Wiley: 1267–1278.
583 Nussbaumer R, Schmid B, Bauer S, et al. (2022) Favorable winds speed up bird migration in spring but
584 not in autumn. *Ecology and evolution* 12(8). Wiley Online Library: e9146.
585 Owen JC and Moore FR (2008) Relationship between energetic condition and indicators of immune
586 function in thrushes during spring migration. *Canadian journal of zoology* 86(7). Canadian Science
587 Publishing: 638–647.
588 Price ER (2010) Dietary lipid composition and avian migratory flight performance: Development of a
589 theoretical framework for avian fat storage. *Comparative biochemistry and physiology. Part A,*
590 *Molecular & integrative physiology* 157(4). Elsevier: 297–309.
591 Prop J, Black JM and Shimmings P (2003) Travel schedules to the high arctic: barnacle geese trade-off
592 the timing of migration with accumulation of fat deposits. *Oikos* 103(2). Wiley: 403–414.
593 Pulgarín-R PC, Gómez C, Bayly NJ, et al. (2019) Migratory birds as vehicles for parasite dispersal?
594 Infection by avian haemosporidians over the year and throughout the range of a long‐distance
595 migrant. *Journal of biogeography* 46(1). Wiley: 83–96.
596 Ramenofsky M (1990) Fat storage and fat metabolism in relation to migration. *Bird migration:*
597 *physiology and ecophysiology*. Springer-Verlag New York: 214–231.
598 Reed KD, Meece JK, Henkel JS, et al. (2003) Birds, migration and emerging zoonoses: west nile virus,
599 lyme disease, influenza A and enteropathogens. *Clinical medicine & research* 1(1): 5–12.
600 Richardson WJ (1978) Timing and Amount of Bird Migration in Relation to Weather: A Review. *Oikos*
601 [30(2). [Nordic Society Oikos, Wiley]: 224–272.](http://paperpile.com/b/9rwraW/NzfN)
602 Ricklefs RE, Medeiros M, Ellis VA, et al. (2017) Avian migration and the distribution of malaria
603 parasites in New World passerine birds. *Journal of biogeography* 44(5). Wiley: 1113–1123.
604 Rotics S, Kaatz M, Turjeman S, et al. (2018) Early arrival at breeding grounds: Causes, costs and a trade-
605 off with overwintering latitude. *The Journal of animal ecology* 87(6): 1627–1638.
606 Rubenstein DR and Hobson KA (2004) From birds to butterflies: animal movement patterns and stable
607 isotopes. *Trends in ecology & evolution* 19(5): 256–263.
608 Saino N, Rubolini D, Jonzén N, et al. (2007) Temperature and rainfall anomalies in Africa predict timing
609 of spring migration in trans-Saharan migratory birds. *Climate Research* 35. Inter-Research Science
610 Center: 123–134.
611 Santiago-Alarcon D, Mettler R, Segelbacher G, et al. (2013) Haemosporidian parasitism in the
612 blackcapSylvia atricapillain relation to spring arrival and body condition. *Journal of avian biology*
613 44(6). Wiley: 521–530.
614 Satterfield DA, Marra PP, Sillett TS, et al. (2018) Responses of migratory species and their pathogens to
615 supplemental feeding. *Philosophical transactions of the Royal Society of London. Series B,*
616 *Biological sciences* 373(1745).
617 Sauer PE, Schimmelmann A, Sessions AL, et al. (2009) Simplified batch equilibration for D/H
618 determination of non-exchangeable hydrogen in solid organic material. *Rapid communications in*
619 *mass spectrometry: RCM* 23(7). Wiley: 949–956.
620 Schimmelmann A (1991) Determination of the concentration and stable isotopic composition of
621 nonexchangeable hydrogen in organic matter. *Analytical chemistry* 63(21). American Chemical
622 Society: 2456–2459.
623 Schimmelmann A, Lewan MD and Wintsch RP (1999) D/H isotope ratios of kerogen, bitumen, oil, and
624 water in hydrous pyrolysis of source rocks containing kerogen types I, II, IIS, and III. *Geochimica et*
625 *cosmochimica acta* 63(22). Elsevier: 3751–3766.
626 Schmaljohann H, Liechti F and Bruderer B (2009) Trans-Sahara migrants select flight altitudes to
627 minimize energy costs rather than water loss. *Behavioral ecology and sociobiology* 63(11). Springer:
628 1609–1619.
629 Schulte-Hostedde AI, Zinner B, Millar JS, et al. (2005) Restitution of mass–size residuals: Validating
630 body condition indices. *Ecology* 86(1). Wiley: 155–163.
631 Serra-Cobo J, Sanz-Trullén V and Martínez-Rica JP (1998) Migratory movements of Miniopterus
632 schreibersii in the north-east of Spain. *Acta theriologica* 43(3). rcin.org.pl: 271–283.
633 Sinelschikova A, Kosarev V, Panov I, et al. (2007) The influence of wind conditions in Europe on the
634 advance in timing of the spring migration of the song thrush (Turdus philomelos) in the south-east
635 Baltic region. *International journal of biometeorology* 51(5). Springer: 431–440.
636 Singh D, Reed SR, Kimmitt AA, et al. (2019) Breeding at higher latitude as measured by stable isotope is
637 associated with higher photoperiod threshold and delayed reproductive development in a songbird.
638 *bioRxiv*. Available at: <https://www.biorxiv.org/content/10.1101/789008> (accessed 22 October 2023).
639 Slowinski SP, Fudickar AM, Hughes AM, et al. (2018) Sedentary songbirds maintain higher prevalence
640 of haemosporidian parasite infections than migratory conspecifics during seasonal sympatry. *PloS*
641 *one* 13(8): e0201563.
642 Smith RJ and Moore FR (2005) Arrival timing and seasonal reproductive performance in a long-distance
643 migratory landbird. *Behavioral ecology and sociobiology* 57(3). Springer Science and Business
644 Media LLC: 231–239.
645 Stott P (2016) How climate change affects extreme weather events. *Science* 352(6293). science.org:

646	1517–1518.

647 Studds CE and Marra PP (2005) Nonbreeding habitat occupancy and population processes: An upgrade
648 experiment with a migratory bird. *Ecology* 86(9). Wiley: 2380–2385.
649 Talbott KM, Becker DJ, Soini HA, et al. (2022) Songbird preen oil odour reflects haemosporidian
650 parasite load. *Animal behaviour* 188. Elsevier: 147–155.
651 Taylor P, Crewe T, Mackenzie S, et al. (2017) The Motus Wildlife Tracking System: a collaborative
652 research network to enhance the understanding of wildlife movement. *Avian Conservation and*
653 *Ecology/Ecologie et Conservation des Oiseaux* 12(1). The Resilience Alliance.
654 Therneau TM and Grambsch PM (2000) *Modeling Survival Data: Extending the Cox Model*. Springer
655 US.
656 Thorsen S (1995-2023) Past Weather in Bloomington, Indiana, USA — March 2020. Available at:
657 <https://www.timeanddate.com/> (accessed 1 November 2022).
658 Tonra CM, Marra PP and Holberton RL (2011) Migration phenology and winter habitat quality are
659 related to circulating androgen in a long-distance migratory bird. *Journal of avian biology* 42(5).
660 Wiley: 397–404.
661 Tøttrup AP, Rainio K, Coppack T, et al. (2010) Local temperature fine-tunes the timing of spring
662 migration in birds. *Integrative and comparative biology* 50(3). academic.oup.com: 293–304.
663 Usui T, Butchart SHM and Phillimore AB (2017) Temporal shifts and temperature sensitivity of avian
664 spring migratory phenology: a phylogenetic meta-analysis. *The Journal of animal ecology* 86(2).
665 Wiley: 250–261.
666 Valkiunas G, Iezhova TA, Krizanauskiene A, et al. (2008) A comparative analysis of microscopy and
667 PCR-based detection methods for blood parasites. *The Journal of parasitology* 94(6): 1395–1401.
668 Van Doren BM and Horton KG (2018) A continental system for forecasting bird migration. *Science*

669	361(6407). science.org: 1115–1118.

670 Visser ME, Perdeck AC, van BALEN JH, et al. (2009) Climate change leads to decreasing bird migration
671 distances. *Global change biology* 15(8). Wiley: 1859–1865.
672 Wanamaker SM, Singh D, Byrd AJ, et al. (2020) Local adaptation from afar: migratory bird populations
673 diverge in the initiation of reproductive timing while wintering in sympatry. *Biology letters* 16(10):
674 20200493.
675 Wang X, Anadón A, Wu Q, et al. (2018) Mechanism of Neonicotinoid Toxicity: Impact on Oxidative
676 Stress and Metabolism. *Annual review of pharmacology and toxicology* 58: 471–507.
677 Weber TP, Houston AI and Ens BJ (1994) Optimal departure fat loads and stopover site use in avian
678 migration: an analytical model. *Proceedings. Biological sciences / The Royal Society* 258(1351). The
679 Royal Society: 29–34.
680 Wikelski M, Tarlow EM, Raim A, et al. (2003) Costs of migration in free-flying songbirds. *Nature*
681 423(6941). Nature Publishing Group: 704–704.
682 Wilcove DS and Wikelski M (2008) Going, going, gone: is animal migration disappearing. *PLoS biology*
683 6(7). journals.plos.org: e188.
684 Witter MS and Cuthill IC (1993) The ecological costs of avian fat storage. *Philosophical transactions of*
685 *the Royal Society of London. Series B, Biological sciences* 340(1291). royalsocietypublishing.org:
686 73–92.
687 Wood SN (2017) *Generalized Additive Models: An Introduction with R, Second Edition*. CRC Press.
688 Wunder MB (2010) Using isoscapes to model probability surfaces for determining geographic origins.
689 *Isoscapes*. Springer. Epub ahead of print 2010.
690 Yorinks N and Atkinson CT (2000) Effects of Malaria on Activity Budgets of Experimentally Infected
691 Juvenile Apapane (Himatione Sanguinea). *The Auk* 117(3). Oxford Academic: 731–738.
692 Zhang J and Wu L (2018) The influence of population movements on the urban relative humidity of
693 Beijing during the Chinese Spring Festival holiday. *Journal of cleaner production* 170. Elsevier:
694 1508–1513.
695 Zhang X, Wan H, Zwiers FW, et al. (2013) Attributing intensification of precipitation extremes to human
696 influence. *Geophysical research letters* 40(19). American Geophysical Union (AGU): 5252–5257.

697

698 **Figures**
699 **Figure 1**. Estimated breeding locations and migratory distances based on junco feather δ^2^H. Orange
700 shading represents overlapping geographic breeding assignments of individuals. Estimates reflect cells
701 within the top 10% highest posterior probability (Bowen et al., 2014; Ma et al., 2020; Wunder, 2010).
702 Points on the map represent the centroid of each estimated breeding polygon, colored by distance to
703 location of capture (star; Bloomington, IN). Lines radiating from the star represent subsequent Motus
704 detections after departure. The seasonal breeding range for *Junco hyemalis(Baillie et al., 2004)* is shown
705 in light grey (the species’ full range map was used to construct geographic assignment maps).

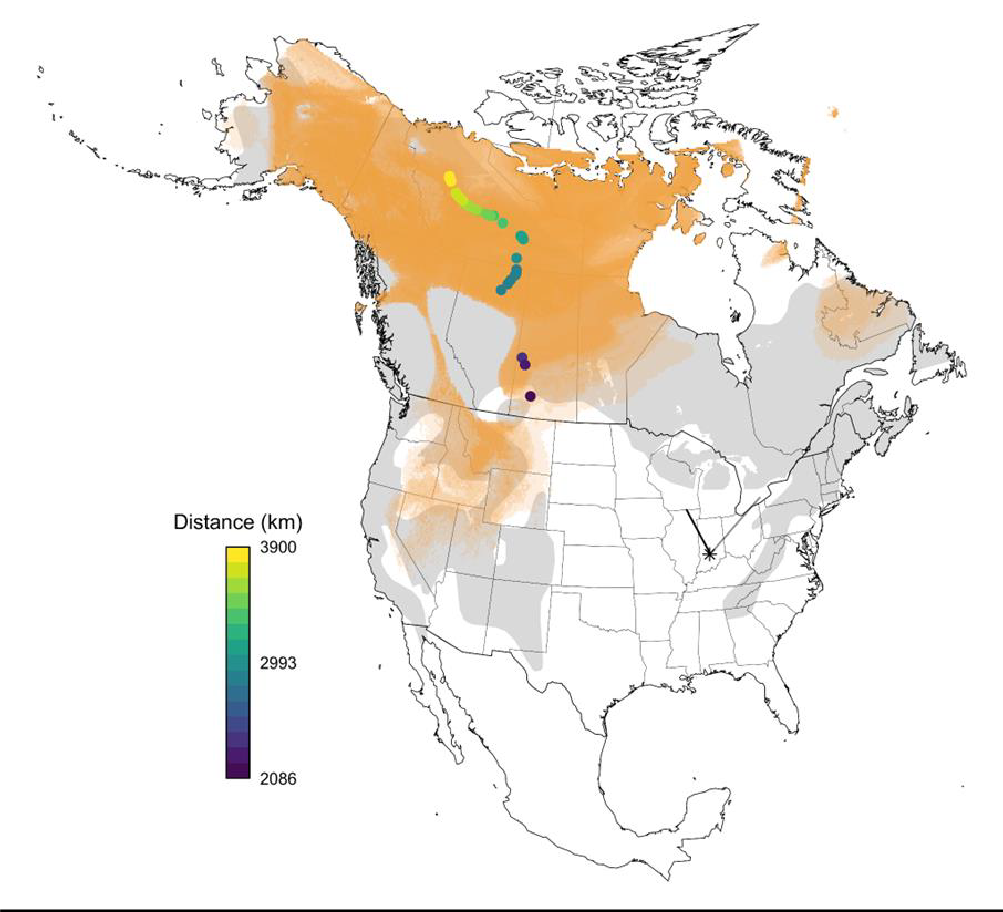
706

707	**Figure 2**. (A) Release and departure timing inferred from Motus at the Kent Farm Research Station for

708	all 37 tagged juncos. (B) Mean probabilities of spring departure as inferred from Kaplan–Meier survival

709

710

711

curves (i.e., 1 – survival probabilities) using the *survival* package alongside 95% confidence intervals.

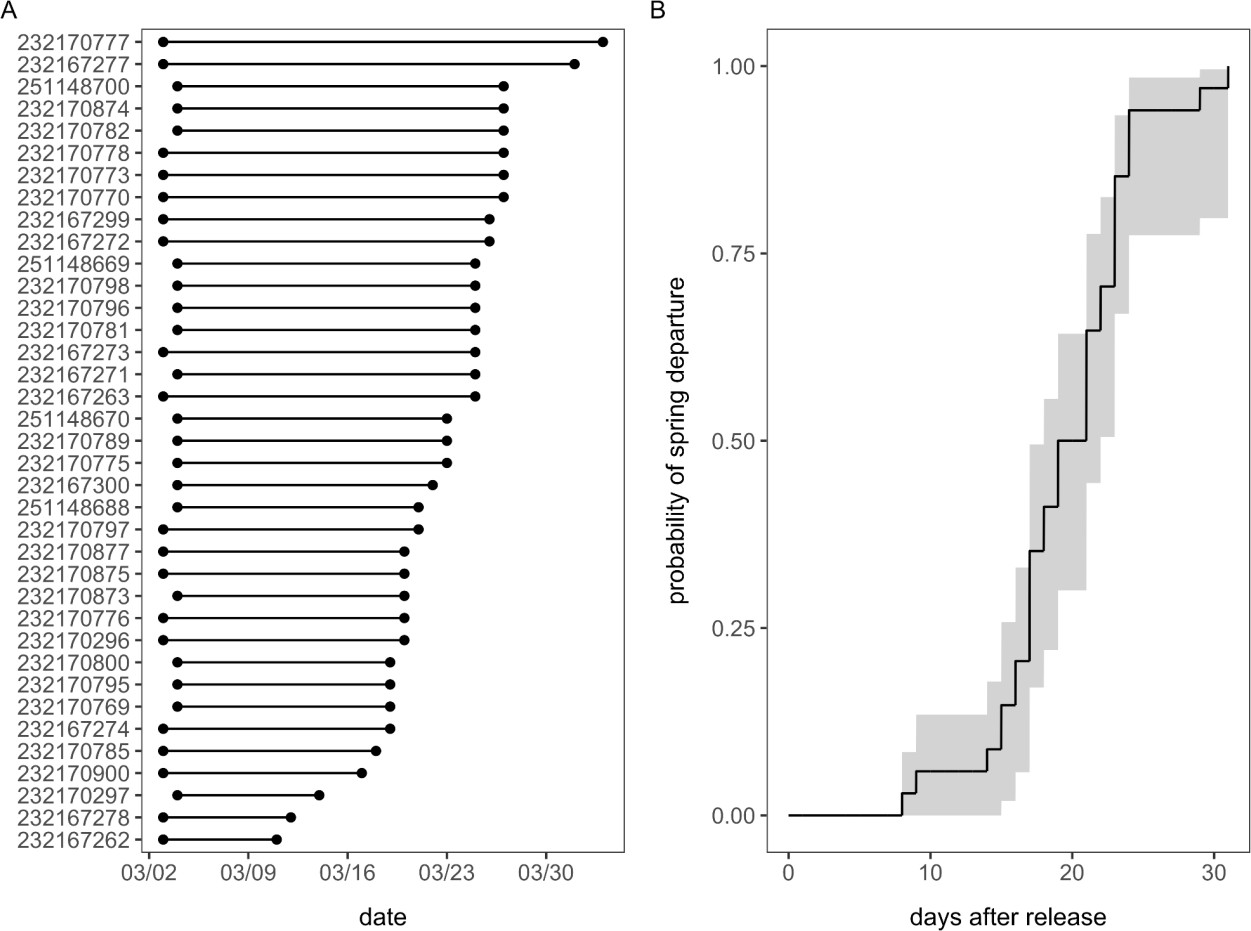

712 **Figure 3.** Relative hazards of spring departure estimated from the most competitive CPHMM as selected
713 through AICc (*w_i_* = 0.61). The predicted relative hazard and 95% confidence intervals from this model are
714 displayed for mean wind direction and weather PC1.
715
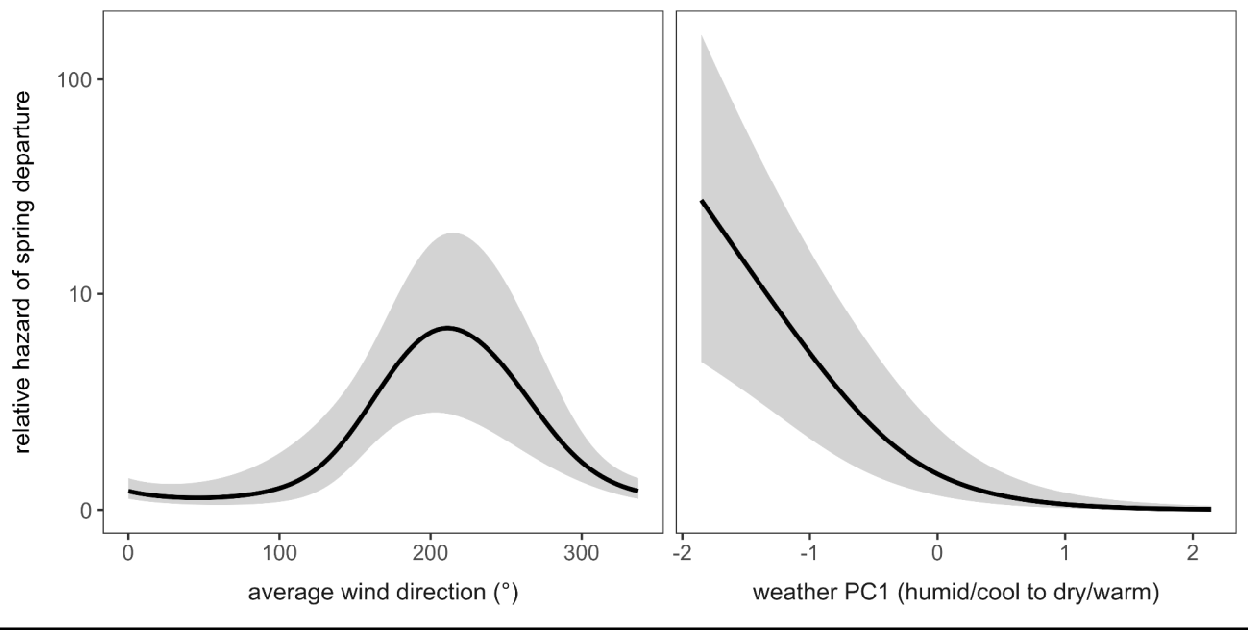

716 **Tables**
717 **Table 1**. Comparison of CPHMMs predicting junco risk of spring departure; all models include a random
718 effect of site. Candidate models are ranked by ΔAICc alongside Akaike weights (*w_i_*) and deviance
719 explained (DE).

| Fixed effects | ΔAICc | *w_i_* | DE |
| --- | --- | --- | --- |
| ~ s(wind direction) + s(weather PC1) | 0.00 | 0.61 | 31% |
| ~ s(wind speed) + s(wind direction) + ti(wind speed, wind direction) | 2.09 | 0.22 | 3% |
| ~ s(wind speed) + s(wind direction) + s(migration distance) | 2.69 | 0.16 | 4% |
| ~ s(intensity) + s(weather PC1) + ti(intensity, weather PC1) | 8.04 | 0.01 | 20% |
| ~ s(intensity) + s(HL ratios) + s(weather PC1) | 12.71 | <0.01 | 10% |
| ~ s(intensity) + s(total leukocytes) + ti(intensity, total leukocytes) | 26.54 | <0.01 | 25% |
| ~ s(body condition) + s(migration distance) | 27.87 | <0.01 | 5% |
| ~ s(fat score) + s(migration distance) | 27.87 | <0.01 | 5% |
| ~ s(intensity) + s(total leukocytes) + s(HL ratios) | 28.15 | <0.01 | 4% |
| ~ s(intensity) + s(HL ratios) + ti(intensity, HL ratios) | 28.15 | <0.01 | 4% |

720
